# *C. elegans* SAS-1 ensures centriole integrity and ciliary function, and operates with SSNA-1

**DOI:** 10.1101/2025.04.22.650004

**Authors:** Keshav Jha, Alexander Woglar, Coralie Busso, Georgios N. Hatzopoulos, Tatiana Favez, Pierre Gönczy

**Affiliations:** Swiss Institute for Experimental Cancer Research (ISREC), School of Life Sciences, Swiss Federal Institute of Technology Lausanne (EPFL), Lausanne, Switzerland

## Abstract

Centrioles are microtubule-based organelles critical for signaling, motility and division. The microtubule-binding protein SAS-1 is homologous to the human ciliopathy component C2CD3 and contributes to centriole integrity in *C. elegans*, but how this function is exerted is incompletely understood. Here, through the generation of a null allele and analysis with U-Ex-STED, we establish that SAS-1 is dispensable for the onset of centriole assembly, but essential for organelle integrity during oogenesis, spermatogenesis and in the early embryo. Additionally, we uncover that SAS-1 is present at the transition zone of sensory neurons, where it contributes in a partially redundant manner to ciliary function. Furthermore, we investigate the relationship between SAS-1 and the *C. elegans* Sjögren Syndrome Nuclear Antigen 1 protein SSNA-1, establishing that SSNA-1 localizes next to the SAS-1 C-terminus in the centriole architecture. Moreover, through molecular epistasis experiments with null alleles of both components, we reveal that SAS-1 is essential for SSNA-1 localization to centrioles during oogenesis and to the transition zone during ciliogenesis. Moreover, using a heterologous human cell assay, we establish that SAS-1 recruits SSNA-1 to microtubules. Overall, our findings help clarify how SAS-1, together with SSNA-1, ensures centriole integrity and reveals that it contributes to cilium function.

## Introduction

Centrioles are 9-fold radially symmetric microtubule-based organelles present across the eukaryotic tree of life (reviewed in Winey & O’Toole 2014; Loncarek & Bettencourt-Dias 2018). Centrioles template the axoneme of cilia and flagella, as well as recruit pericentriolar material (PCM) to form centrosomes, which act as microtubule organizing centers (MTOCs) in animal cells. As a result of these fundamental roles, centrioles are critical for signaling, motility and division. Cells maintain tight control over centriole number and architecture, and failure to do so can result in disease (reviewed in Braun & Hildebrandt 2017; Nigg & Holland 2018). In most proliferating cells, two pre-existing centrioles seed the formation of one procentriole each in their vicinity. Centrioles are extremely stable in general, but undergo programmed elimination in some circumstances, including during oogenesis of metazoan species (reviewed in Manandhar et al. 2005; Kalbfuss & Gönczy 2023b). Overall, despite their importance, the mechanisms regulating centriole integrity remain incompletely understood.

Systematic analyses in *C. elegans* identified a core set of evolutionarily conserved components necessary for centriole assembly, comprising SAS-7, SPD-2, ZYG-1, SAS-6, SAS-5 and SAS-4 (reviewed in Ohta et al. 2017; Pintard & Bowerman 2019). In brief, centriole assembly begins when the kinase ZYG-1 phosphorylates SAS-5 on a platform formed by SAS-7 and SPD-2. Phosphorylated SAS-5 then associates with SAS-6, which can self-assemble into an inner tube element thought to scaffold recruitment of the microtubule-binding protein SAS-4 and then of centriolar microtubules. *C. elegans* centrioles are merely ∼150 nm by ∼120 nm in dimensions (Sugioka et al. 2017; Woglar et al. 2022), yet the distribution of centriolar and PCM components has been determined with high accuracy using Ultra-Expansion coupled to STED super-resolution microscopy (U-Ex-STED) (Woglar et al. 2022).

Although *C. elegans* lacks motile cilia or flagella, centrioles seed the ciliary axoneme of 60 sensory neurons, 56 in the head region and 4 in the tail region of the animal (Inglis 2006; Ward et al. 1975). After templating ciliary axoneme formation, centrioles degenerate and are thus absent from mature cilia (Serwas et al. 2017; Nechipurenko et al. 2017). By contrast, PCM proteins such as SPD-5 and TBG-1 (γ- tubulin) remain at the ciliary base, where they function as acentriolar MTOCs (Magescas et al. 2021; Garbrecht et al. 2021). Distal to SPD-5 and TBG-1 lies the transition zone, which marks the beginning of the cilium proper and comprises a central cylinder positioned internal to axonemal microtubules, as well as Y-links that bridge the outside of axonemal microtubules with the ciliary membrane (Inglis 2006; Ward et al. 1975).

Centrioles are also eliminated in other physiological contexts in *C. elegans*. Thus, centrioles undergo programmed elimination in multiple lineages during embryogenesis, resulting in merely 68 out of 558 cells retaining centrioles by the first larval stage (L1) (Kalbfuss & Gönczy 2023a). Moreover, as in other metazoan organisms, centrioles are eliminated during oogenesis, with sperm contributing the sole pair of centrioles to the zygote, thereby ensuring bipolar spindle assembly and faithful chromosome segregation (Mikeladze-Dvali et al. 2012; Pierron et al. 2023). Correlative Light Electron Microscopy (CLEM) analysis revealed that oogenesis centriole elimination begins during meiotic prophase with the loss of the central tube element located inside the wall of centriolar microtubules (Pierron et al. 2023). Whereas in Drosophila removal of the kinase Polo, and thereby of the PCM, is critical for triggering oogenesis centriole elimination (Pimenta-Marques et al. 2016), the related Polo-like kinases and the PCM do not modulate this process in *C. elegans* (Pierron et al. 2023).

On the other hand, SAS-1 plays an important role in regulating oogenesis centriole elimination in *C. elegans* (Pierron et al. 2023). SAS-1 is a C2-domain containing protein homologous to human C2CD3, which is essential for assembly of the centriole distal segment, with patient mutations leading to an oral-facial-digital (OFD) syndrome with severe microcephaly and cerebral malformations (Hoover et al. 2008; Balestra et al. 2013; Thauvin-Robinet et al. 2014). SAS-1 localizes to the central tube of *C. elegans* centrioles, with the N-terminus located next to centriolar microtubules and the C-terminus positioned more centrally (Woglar et al. 2022). When expressed in human cells, *C. elegans* SAS-1 binds to and stabilizes microtubules (Von Tobel et al. 2014). Importantly, during oogenesis in *C. elegans*, SAS-1 is the first known protein to leave centrioles, followed by the loss of organized centriolar microtubules and removal of other centriolar proteins, including SAS-4 and SAS-6 (Pierron et al. 2023). In the reduction of function allele *sas-1*(*t1521*), centrioles are eliminated precociously during oogenesis (Pierron et al. 2023). Moreover, centrioles contributed to the zygote by *sas-1*(*t1476*) or *sas-1*(*t1521*) mutant sperm lose integrity shortly after fertilization (Von Tobel et al. 2014). Furthermore, compromising maternal *sas-1* function using these mutant alleles also results in centriole instability, in this case during the course of embryogenesis (Von Tobel et al. 2014). Whether a null allele of *sas-1* would result in a more severe phenotype, perhaps complete failure of centriole assembly, remains to be determined.

Here, we generated and analyzed a null allele of *sas-1*, thereby establishing that the protein is dispensable for centriole assembly, but critical for organelle integrity. We also report that SAS-1 localizes to the transition zone of sensory cilia and contributes to their function. Furthermore, we investigated the relationship of SAS-1 with SSNA-1, the worm homologue of Sjögren Syndrome Nuclear Antigen 1 protein SSNA-1, finding that SAS-1 is essential for recruiting SSNA-1 to centrioles and cilia in *C. elegans*, as well as to microtubules in a heterologous assay. Together, our work helps to understand how SAS-1 and SSNA-1 operate to ensure integrity of the centriole organelle.

## RESULTS

### Loss of SAS-1 leads to premature centriole elimination during oogenesis

Previous analyses of SAS-1 were conducted with the reduction of function alleles *sas-1*(*t1476*) and *sas-1*(*t1521*) (Gönczy et al. 1999; Von Tobel et al. 2014; Pierron et al. 2023). To investigate the consequences of complete loss of SAS-1, we generated a null allele using CRISPR/Cas9 (Fig. S1A). The resulting *sas-1(is13*) mutant harbors a deletion spanning the 5’-UTR and part of exon 1, which removes the ATG start codon and generates a protein null (Fig. S1B-D). Heterozygous *sas-1(is13*)/hT2 worms give rise to ∼20% *sas-1(is13*) homozygous animals at all tested temperatures (Fig. S1E). However, such homozygous animals lay very few embryos, which all fail to hatch, indicating that *sas-1* exerts an essential function.

Equipped with the *sas-1(is13*) null allele, we set out to investigate the consequences of SAS-1 loss, first by analyzing proliferating germ cell nuclei and associated centrioles in the mitotic region of the gonad using immunofluorescence (Fig. 1A-D). In control worms, nuclei in this region are similarly sized, with each nucleus associated with centrioles marked by SAS-7 and SAS-4 (Fig. 1B, 1E). By contrast, *sas-6(ok2554)* null mutant animals harbor much fewer nuclei in this region (Fig. 1F) (Mikeladze-Dvali et al. 2012). Moreover, *sas-6(ok2554)* mutant nuclei are often much larger than normal (Fig. 1E), and almost never associated with a centriolar focus (Fig. 1C; 3.2%, N = 348), together reflecting failed chromosome segregation and lack of centriole assembly. In *sas-1(is13)* null mutants, we found that most nuclei are similarly sized (Fig. 1D, 1E), and usually associated with centrioles marked by SAS-7 and SAS-4, as in the control (Fig. 1D, 1F; 79.2%, N = 645). The fraction of nuclei exhibiting such association remains constant as germ cell nuclei mature in *sas-1(is13)* animals (Fig. S2A-C). Furthermore, live imaging of the mitotic region of the gonad in control and *sas-1(is13)* mutant animals revealed that SAS-4::GFP levels are unchanged compared to the control (Fig. S2D-F), whereas those of GFP::SAS-7 are lower (Fig. S2G-I). Overall, we conclude that centrioles can assemble and recruit SAS-7 and SAS-4 despite the absence of SAS-1, although centrioles are lacking next to some nuclei.

**Figure 1.**
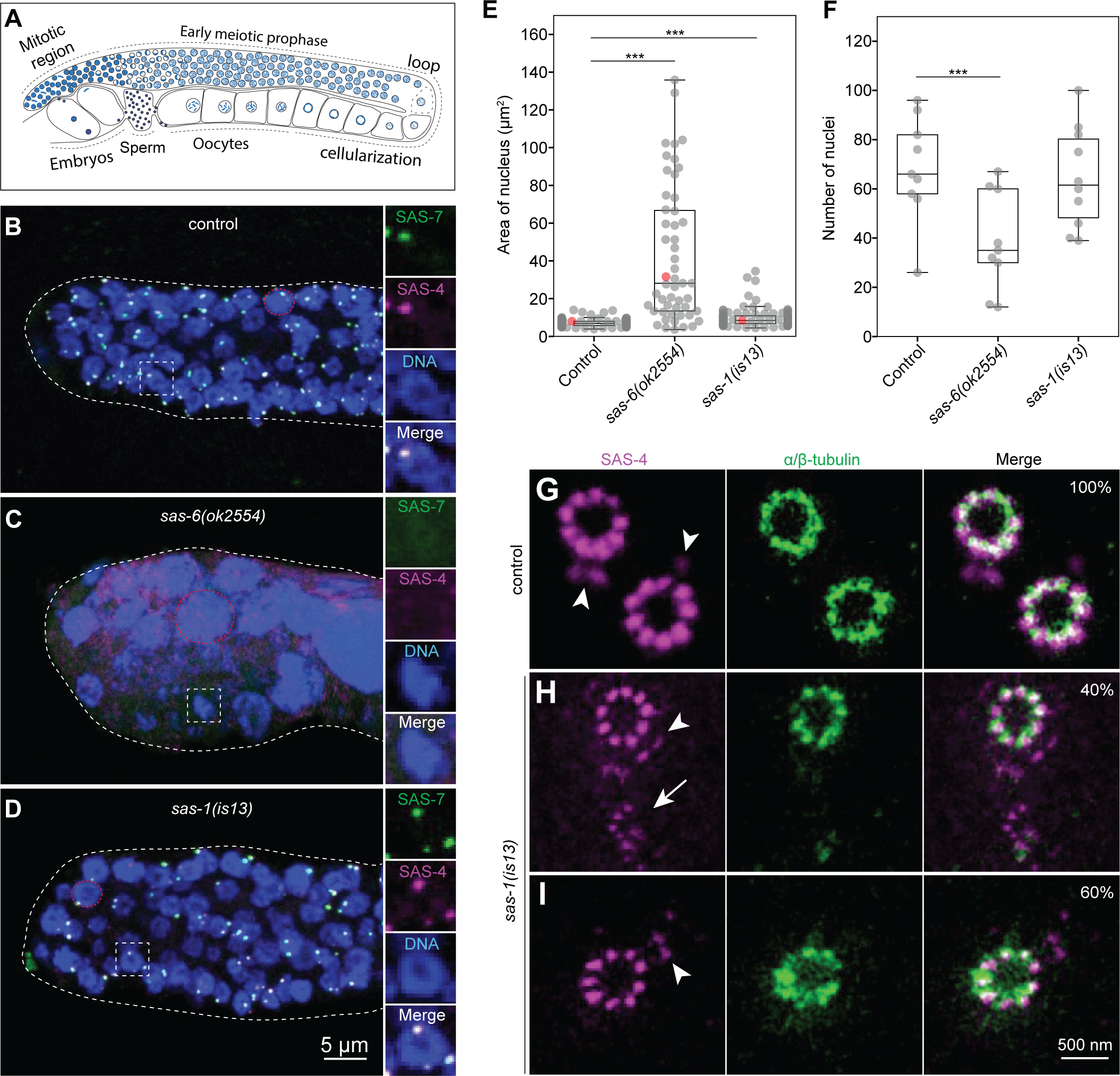
SAS-1 loss leads to premature centriole elimination during oogenesis. **A.** Schematic of the gonad of a young adult hermaphrodite showing different stages of gametogenesis, with the early mitotic region, followed by the meiotic prophase, before the loop region and cellularization. Mature oocytes traverse the spermatheca where fertilization takes places. **B-D.** Representative immunofluorescence images of the early mitotic region in fixed gonads of control (B), *sas-6(ok2554)* (C), and *sas-1(is13)* (D) animals expressing GFP::SAS-7, immunostained with antibodies against SAS-4 and GFP. Dotted lines highlight edges of gonad. Square insets on the right are ∼1.5 times magnified, whereas dashed red circles exemplify regions measured for nuclear area reported in (E). Brightness and contrast adjusted individually for each image. **E.** Quantification of nuclear area in the mitotic region of indicated genotypes. Number of nuclei: N *=* 98 (control, from 5 gonads); 51 (*sas-6(ok2554)*, from 6 gonads); 101 (*sas-1(is13)*, from 5 gonads). Student’s two-tailed t-tests, whereby *P* < 0.001 (***). **F.** Number of nuclei in the mitotic region of gonads of indicated genotypes. Number of gonads *=* 9 (control and *sas-6(ok2554);* 10 (*sas-1(is13)*). Student’s two-tailed t-tests, whereby *P* < 0.01 (**). The difference between control and *sas-1*(*is13*) is not significant (*P* = 0.68). **G-I.** Representative U-Ex-STED images of centrioles in the mitotic region of control (G; N = 23) and *sas-1(is13)* mutants (H, I; N = 15), immunostained for SAS-4 and α/β-tubulin. Arrowheads: procentrioles; arrow: dismantled centriole.

We next report high spatial resolution analysis of centriole architecture in the mitotic region of the gonad using U-Ex-STED (Gambarotto et al. 2019; Woglar et al. 2022). In the wild-type, the two mature centrioles are recognized in cross-sectional views by the radial distribution of SAS-4 and microtubules (Fig. 1G). In addition, SAS-4 is present in the two immature procentrioles at much reduced levels (Fig. 1G, arrowheads) (Woglar et al. 2022). We observed that signal intensities for SAS-4 and microtubules are weaker in the *sas-1(is13)* null mutant compared to the control (Fig. 1H, 1I, compare to Fig. 1G). Moreover, centrioles are slightly narrower in *sas-1(is13)* (Fig. S2J). Since SAS-4::GFP levels are unchanged in the mitotic region of the gonad (see Fig. S2D-F), this observation suggests that, albeit being assembled in the first place, centrioles lacking SAS-1 have compromised integrity, a fragility potentially compounded by the expansion protocol. In addition, we noted two centriole architecture configurations in the mitotic region of the gonad of *sas-1(is13)* mutants: in 40% of cases, one of the two centrioles is deformed and barely visible (Fig. 1H, arrow), whereas in the remaining ∼60% only one centriole is detected (Fig. 1I). In both configurations, next to the mature centriole lies one procentriole, distinguished by its weak levels of SAS-4 (Fig. 1H, 1I; arrowheads). We speculate that the more intact centrioles are younger than the ones that are deformed or absent, reflecting loss of organelle integrity over time.

Overall, these observations indicate that centrioles can assemble during oogenesis in the absence of SAS-1, but that their integrity is compromised.

### SAS-1 is dispensable for centriole assembly but essential for organelle integrity in the embryo and sperm cells

We sought to test further whether SAS-1 is dispensable for the onset of centriole assembly, since in principle centriole assembly in the mitotic region of the gonad could have been driven by residual maternally-supplied SAS-1 persisting from the heterozygous parent. To this end, we conducted marked mating experiments, in which otherwise wild-type males expressing TagRFP::SAS-7 are mated with *sas-1(is13)* mutant hermaphrodites expressing SAS-4::GFP (Fig. S3). If SAS-1 is essential for centriole assembly, then SAS-4::GFP should not be recruited to centrioles in the resulting one-cell stage embryos; moreover, a monopolar spindle is expected to assemble in each blastomere at the two-cell stage, as in embryos lacking maternal *zyg-1* or *sas-5* function (O’Connell et al. 2001; Delattre et al. 2004) (Fig. S3). Contrary to these predictions, time-lapse microscopy of such embryos established that two foci of SAS-4::GFP are invariably observed at the one-cell stage (Fig. 2A, 0 min; N = 9), and bipolar spindle assembly always occurs in both blastomeres at the two-cell stage (Fig. 2A, 9 min). Interestingly, in addition, we observed that one of the two SAS-4::GFP foci present in each blastomere at the two-cell stage exhibits a weaker signal than the other one initially (Fig. 2A, 6 min, insets 3 and 4), and often becomes undetectable by the end of the cell cycle (Fig. 2A, 9 min, insets 3 and 4, N = 15/18 centrioles). We reason that the initial presence of the centriole must have sufficed to recruit the PCM, since bipolar spindle assembly systematically takes place in both blastomeres at the two-cell stage in such embryos (Fig. 2A, 9 min; N = 9). Overall, these findings further demonstrate that lacking SAS-1 entirely does not prevent centriole assembly, but compromises organelle integrity.

**Figure 2.**
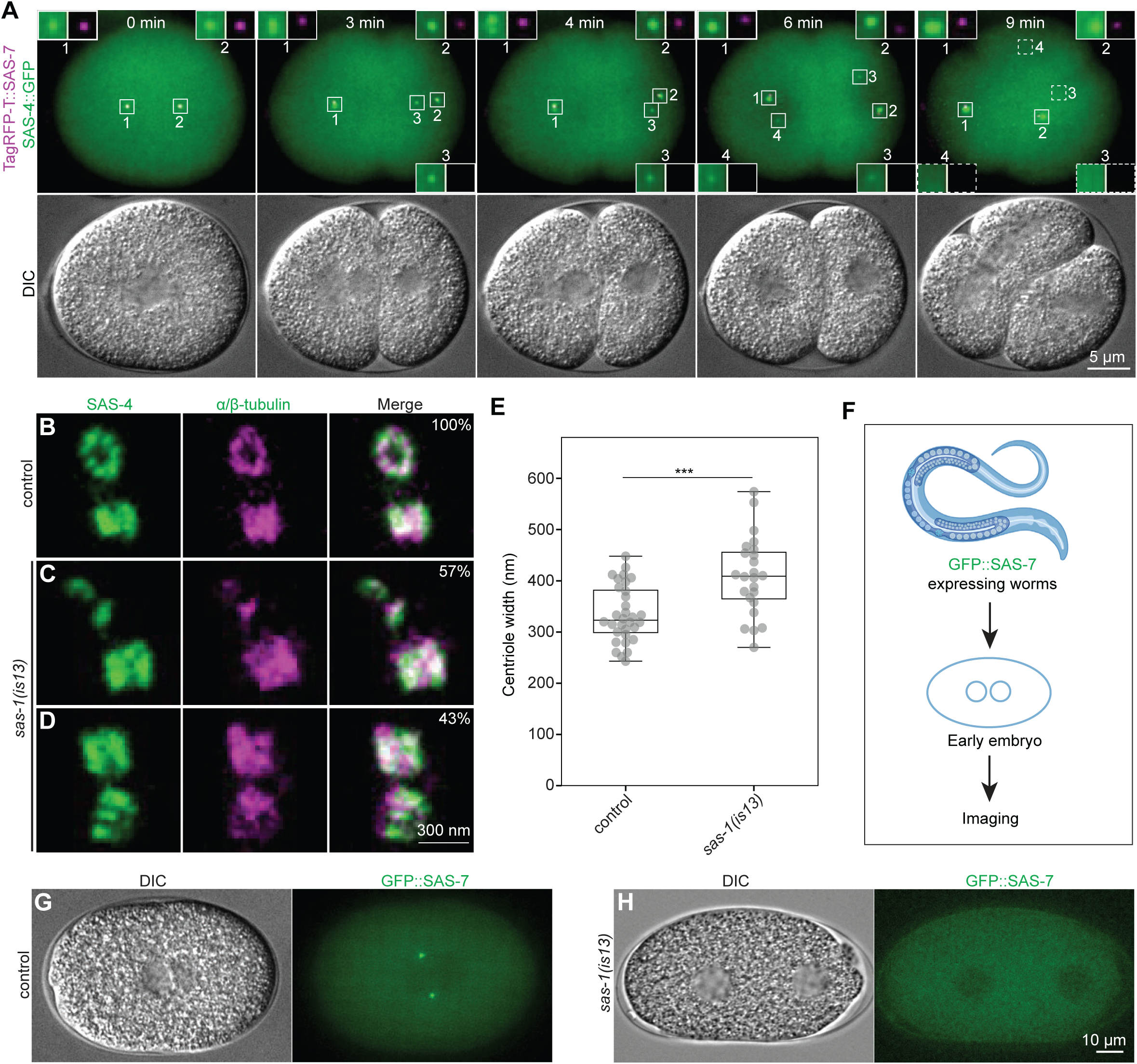
SAS-1 is dispensable for centriole assembly but essential for organelle integrity. **A.** Snapshots from live imaging of early embryo stemming from mating males expressing TagRFP-T::SAS-7 with *sas-1(is13)* mutant hermaphrodite expressing SAS-4::GFP (see Fig. S3). TagRFP-T::SAS-7 marks sperm-contributed centrioles (insets 1 and 2), whereas SAS-4::GFP marks all centrioles assembled after fertilization. T = 0 corresponds to metaphase. Insets 3 and 4 mark degenerating centrioles assembled next to sperm-contributed centrioles. The DIC image is a single plane, whereas the two merged fluorescent channels are maximum intensity projection of selected Z-planes. **B-D.** Representative U-Ex-STED images of sperm centrioles in control (B; N = 16) and *sas-1(is13)* mutants (C, D; N = 23), immunostained for SAS-4 and α/β-tubulin. Note that U-Ex-STED of sperm centrioles requires methanol fixation (unlike that of gonad centrioles in Fig. 1G-I), compromising resolution. **E.** Quantification of sperm centriole width, excluding centriole fragments, in (B-D). N *=* 32 (control), 24 (*sas-1(is13)*). Student’s two-tailed t-tests, whereby *P* < 0.001 (***). **F.** Schematic of live imaging of one-cell stage embryos in G and H (drawn with Biorender). **G, H.** Snapshots from live imaging at the time of pronuclear meeting in control (G; N = 10) and *sas-1(is13)* mutant (H; N = 9) one-cell stage embryos. Images were acquired every minute. The DIC channel is a single plane, whereas the GFP::SAS-7 channel is a maximum intensity projection of selected Z-planes.

We analyzed also the impact of complete loss of SAS-1 in sperm cells. In previous work, electron-microscopy (EM) analysis did not reveal an apparent difference between wild-type and *sas-1(t1476)* sperm centrioles, although mutant sperm-contributed centrioles were then rapidly eliminated in the embryo (Von Tobel et al. 2014). Here, we used U-Ex-STED to examine the molecular architecture of sperm centrioles. In the control, both SAS-4 and microtubules localize to sperm centrioles, as anticipated (Fig. 2B). Moreover, this analysis revealed defects in *sas-1(is13)* mutant sperm centrioles, including increased width and partial disassembly (Fig. 2C-E). What is the fate of these centrioles following fertilization? To address this question, we used live imaging to monitor sperm-contributed centrioles marked by GFP::SAS-7 (Fig. 2F). In the control, both sperm-contributed centrioles are visible at pronuclear meeting (Fig. 2G). By contrast, no GFP::SAS-7 focus is present at the equivalent stage in *sas-1(is13)* mutant embryos (Fig. 2H, N = 9), although one centriole could sometimes be detected earlier in the cell cycle (3/9). Together, these observations indicate that the loss of SAS-1 allows sperm centrioles to be assembled, but also compromises their integrity, presumably leading to their rapid demise in the embryo following fertilization.

### SAS-1 kinetics during embryogenesis

We set out to investigate SAS-1 distribution during programmed centriole elimination in embryogenesis (Kalbfuss & Gönczy 2023a). Embryos complete their proliferative phase by ∼350 minutes post-fertilization (mpf), by which time most of the 558 cells present at hatching have been generated. In the subsequent morphogenesis stage, embryos sequentially adopt characteristic bean, comma, 1.5-fold, and 2-fold configurations, accompanied by extensive lineage-specific centriole elimination (Kalbfuss & Gönczy 2023a). We performed lattice light sheet live imaging of embryos expressing TagRFP-T::SAS-7 and GFP::SAS-1 from the bean stage (∼360 mpf) through the 2-fold stage (∼460 mpf) (Fig. 3A, Movie S1), after which twitching prevents faithful centriole monitoring. We found that the number of GFP::SAS-1 foci decrease from ∼550 (± 28) at the bean stage to ∼474 (± 43) at the 2-fold stage (Fig. 3B). This final number is higher than the ∼220 (± 50) determined previously at this stage for GFP::SAS-7 (Kalbfuss & Gönczy 2023a). Accordingly, some GFP::SAS-1 foci that do not appear to bear TagRFP-T::SAS-7 are present at the 2-fold stage, in particular in the embryo anterior, although this may reflect in part weaker signal intensity for TagRFP-T::SAS-7 (Fig. 3A, 2-fold stage, inset). Moreover, whether these GFP::SAS-1 positive foci correspond to *bona fide* centrioles or merely to a remaining focus of SAS-1 protein remains to be determined.

**Figure 3.**
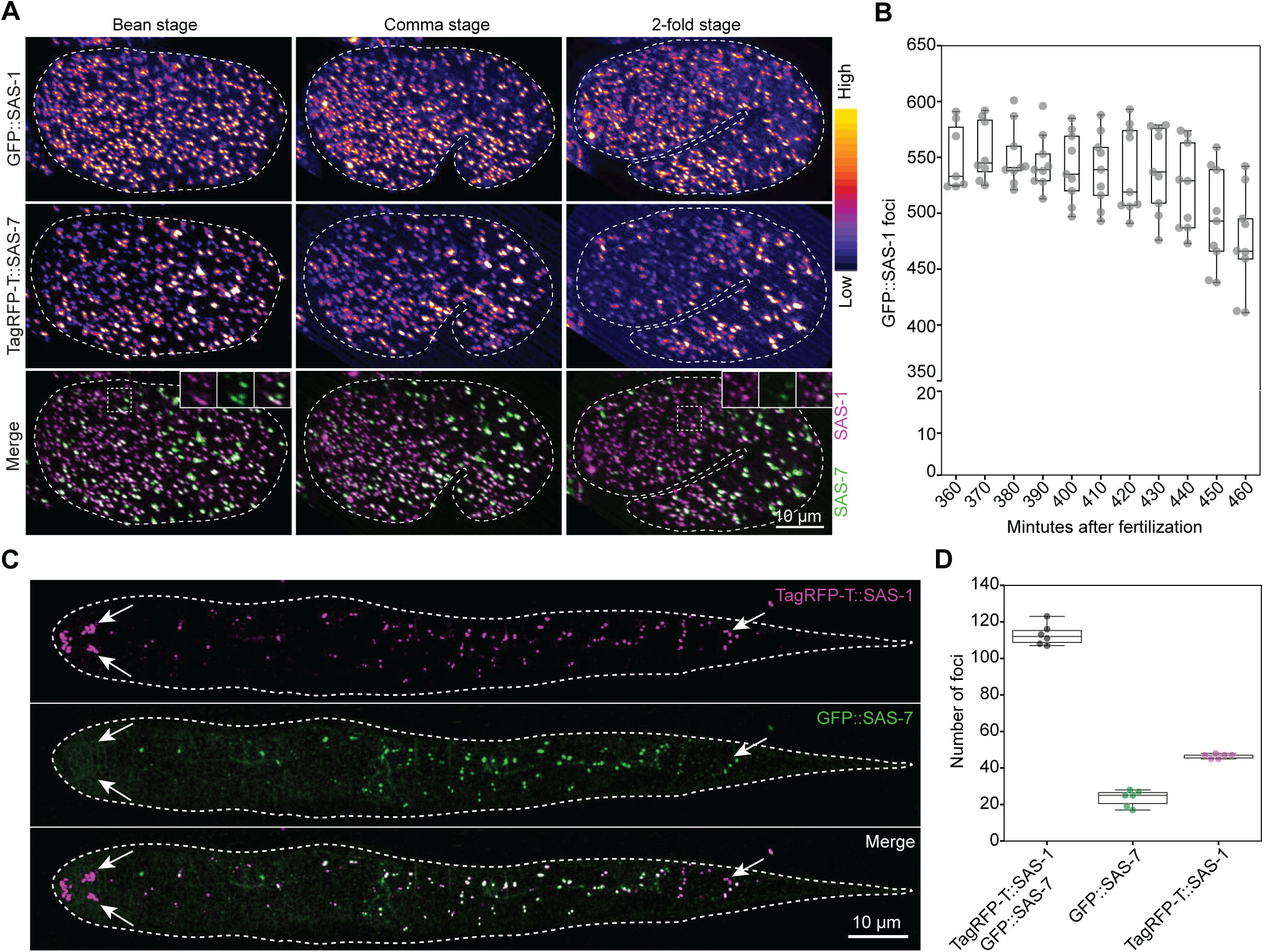
SAS-1 departs from centrioles during embryogenesis programmed centriole elimination. **A.** Snapshots of time-lapse of embryos expressing TagRFP-T::SAS-7 and GFP::SAS-1 at bean, comma, and 2-fold stage. Top and middle rows show single channels displayed with a "fire" LUT, the bottom row the merged of the two channels, with insets highlighting foci in the embryo anterior. See also Movie S1. **B.** Quantification of number of foci positive for TagRFP-T::SAS-1 from the bean stage until the 2-fold stage, 10 min before twitching (N = 9). **C.** Live imaging of L1 larva expressing TagRFP-T::SAS-1 and GFP::SAS-7. Anterior is to the left. Arrows point to foci strongly positive solely for TagRFP-T::SAS-1 in the anterior or posterior of the animal. **F.** Quantification of number of foci positive for TagRFP-T::SAS-1 and GFP::SAS-7, or for just one of the two fusion proteins (N = 6 L1 larvae).

Regardless, despite this difference between the two markers at the 2-fold stage, by the end of embryogenesis, in L1 larvae, TagRFP-T::SAS-1 is present essentially in the same cells as GFP::SAS-7, with a few additional foci positive for only one of the two fusion proteins, typically at low signal intensity, suggestive of spurious labelling (Fig. 3C, 3D). We noted a striking exception to this general colocalization: the presence of strong bilateral TagRFP-T::SAS-1 foci in both anterior and posterior of the animal that do not colocalize with GFP::SAS-7 (Fig. 3C, arrows). Based on their position, these extra TagRFP-T::SAS-1 foci appear to mark the base of anterior and posterior sensory ciliated neurons.

### SAS-1 localizes at the transition zone of sensory cilia

We set out to address whether SAS-1 truly localizes to the ciliary base and, if so, where exactly compared to previously mapped proteins. Live imaging of L1 larvae revealed a consistent separation between TagRFP-T::SAS-1 and GFP::SPD-5 foci, with TagRFP-T::SAS-1 foci being positioned slightly more anterior to GFP::SPD-5 foci in anterior cilia (Fig. 4A), and slightly more posterior to GFP::SPD-5 foci in posterior cilia (Fig. 4B). These patterns suggest that SAS-1 does not localize at the ciliary base proper, but instead in the transition zone just above. Accordingly, we found that TagRFP-T::SAS-1 fully colocalizes with the transition zone protein GFP::MKSR-2 (Bialas et al. 2009), both in anterior and posterior cilia (Fig. 4C, 4D). We next investigated whether SAS-1 is present within the confines of axonemal microtubules, where the central cylinder is located, or instead external from them, where Y-links are located. To mark axonemal microtubules, we placed L1-L2 larvae expressing TagRFP-T::SAS-1 in a solution containing the microtubule probe SPY650-tubulin, reasoning that this small molecule could be taken up by sensory cilia. As shown in Figure 4E, we found this to be the case. Importantly, AiryScan imaging revealed that TagRFP-T::SAS-1 is present within the confines of the axonemal microtubule signal marked by SPY650-tubulin (Fig. 4E, 4F), echoing SAS-1 localization at the central tube inside the centriolar microtubule wall (Woglar et al. 2022).

**Figure 4.**
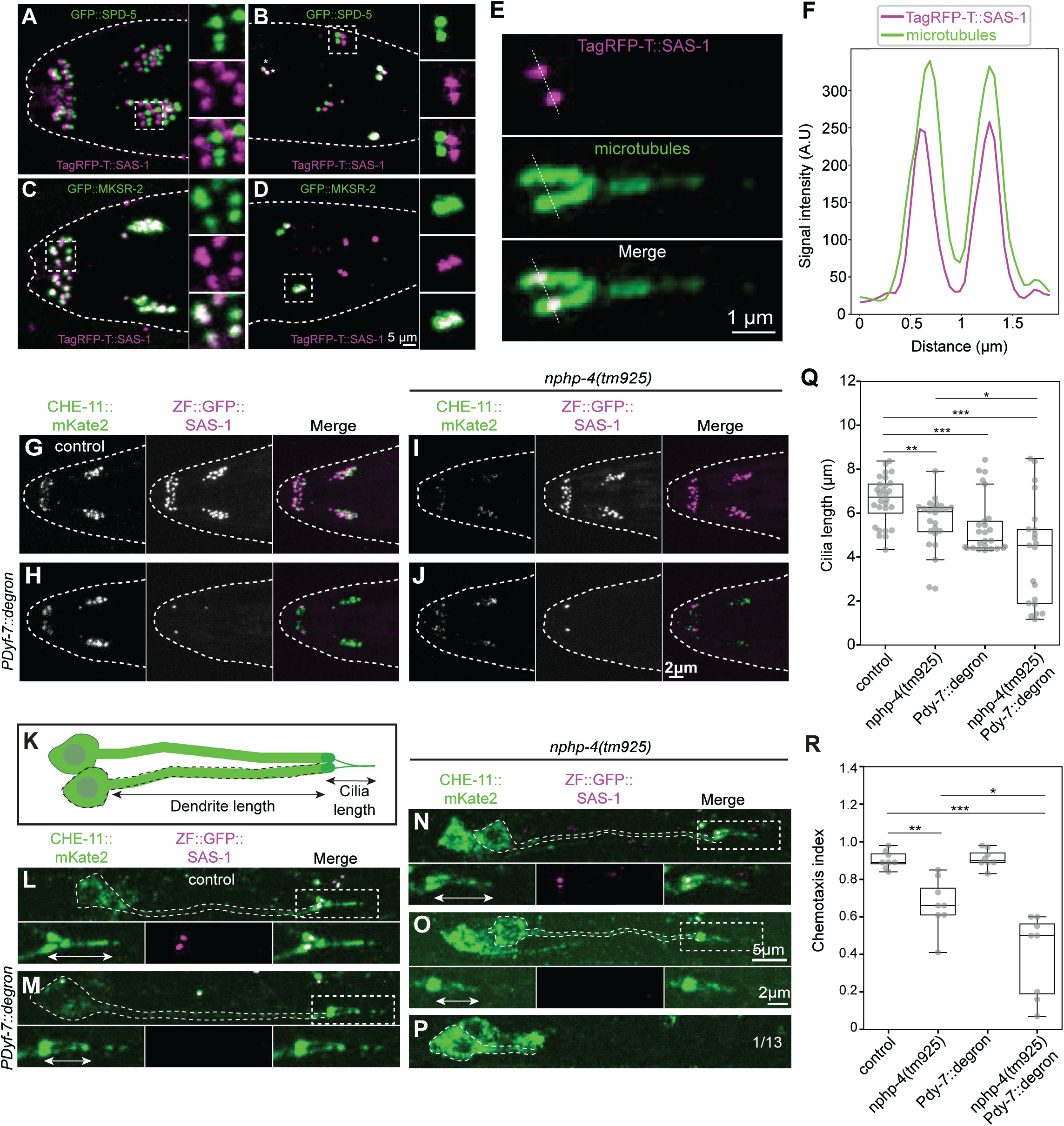
SAS-1 is a component of the transition zone and contributes to ciliary function. **A, B.** Live imaging of anterior (A) and posterior (B) cilia of L1 larva expressing GFP::SPD-5 and TagRFP-T::SAS-1. Insets are magnified approximately two-fold. **C, D.** Live imaging of anterior (C) and posterior (D) cilia of L1 larva expressing GFP::MKSR-2 and TagRFP-T::SAS-1. Insets are magnified approximately two-fold. MKSR-2 localizes exclusively to the transition zone and is absent from centrioles. **E.** Live Airyscan imaging of posterior cilium of L1 larva expressing TagRFP-T::SAS-1, with axonemal microtubules stained with SPY650-tubulin. **F.** Representative line profile of signal intensity of TagRFP-T::SAS-1 and SPY650-tubulin; N = 6 L1 larvae. **G-J.** Live imaging of L1 larvae expressing CHE-11::mKate2 and ZF::GFP::SAS-1 in control (G; N *=* 7), *Pdyf-7::degron* (H; N *=* 10, note two remaining ZF::GFP::SAS-1 foci, likely due to lack of ZIF-1 expression in these neurons), *nphp-4(tm925)* (I; N *=* 6), and *Pdyf-7::degron nphp-4(tm925)* (J; N *=* 10). **K.** Schematic of posterior sensory neuron, with an indication of features monitored in L-P. **L-P.** Live imaging of young adult worms expressing CHE-11::mKate2 and ZF::GFP::SAS-1 in control (L; N *=* 12), *Pdyf-7::degron* (M; N *=* 11), *nphp-4(tm925)* (N; N *=* 13), and *Pdyf-7::degron nphp-4(tm925)* (O, P; N *=* 13). Insets are 1.5-fold magnified. White line: cilia, from the base to the tip of axoneme. Dotted lines: edges of neurons. **Q.** Quantification of cilium length, as schematized in (K). N *=* 28 (control), 25 (*Pdyf-7::degron*), 24 (*nphp-4(tm925)*), 21 (*Pdyf-7::degron nphp-4(tm925)*). Student’s two-tailed t-tests, whereby *P* < 0.05 (*); *P* < 0.01 (**); *P* < 0.001 (***). **R.** Chemotaxis index for animals of indicated genotypes. Data points are from eight experiments. Student’s two-tailed t-tests, whereby *P* < 0.05 (*); *P* < 0.01 (**); *P* < 0.001 (***).

We set out to test whether SAS-1 exerts a function at sensory cilia. We found that the TagRFP-T::SAS-1 signal at the transition zone declines during the larval stages (Fig. S4A), suggesting that a putative function would be exerted during early ciliogenesis. To remove SAS-1 from developing cilia, given that *sas-1(is13)* mutant embryos fail to develop (see Fig. 2H), we deployed ZIF-1-mediated degradation of endogenous SAS-1 using the neuron-specific *dyf-7* promoter, which expresses during early ciliogenesis (Fig. S4B) (Wang, Tang, et al. 2017; Cheerambathur et al. 2019). We found that ZIF-1-mediated depletion of SAS-1 does not decrease levels of the axonemal motor protein CHE-11::mKate2 from anterior cilia (Fig. 4G, 4H). Because transition zone proteins can act in a redundant fashion (Williams et al. 2008; Williams et al. 2011; Schouteden et al. 2015), we tested whether the impact on CHE-11::mKate2 distribution of lacking the Y-link transition zone protein NPHP-4 might be compounded by depleting SAS-1 (Jauregui & Barr 2005; Williams et al. 2010), but found this not to be the case (Fig. 4I, 4J). Nevertheless, we found that posterior cilia length is reduced upon SAS-1 depletion alone, and that this phenotype is exacerbated by combining SAS-1 removal with the null mutant *nphp-4(tm925)* (Fig. 4K-Q). However, we found that dendrite length is largely unaffected (Fig. S4C), with the exception of one instance of complete retrograde dendrite extension failure (Fig. 4P). Next, we conducted a chemosensation assay to determine whether ciliary function is perturbed upon SAS-1 removal. In this assay, worms are placed in the center of a plate that contains a drop of the attractant Isoamyl alcohol on one side and a drop of the repellent Ethanol on the other (Fig. S4D) (Bargmann et al. 1993). Wild-type animals move towards the attractant, whereas animals with defective chemosensation fail to do so, a preference quantified with a chemotaxis index. As shown in Figure 4R, we found that ZIF-1-mediated depletion of SAS-1 from cilia results in a slight but significant chemosensation defect, which is enhanced in the *nphp-4(tm925)* background. Taken together, these findings establish SAS-1 as a transition zone protein that contributes to proper ciliary function in *C. elegans*.

### SSNA-1 colocalizes with SAS-1 at the central tube of centrioles

Prompted by the apparent phenotypic similarity between embryos depleted of maternal SAS-1 (Von Tobel et al. 2014) and those lacking the Sjögren Syndrome Nuclear Antigen 1 (SSNA-1) protein (Pfister et al. 2024), we set out to investigate a possible relationship between these two components. Live imaging of control (Fig. 5A) and *ssna-1(bs182)* null mutant (Fig. 5B) embryos confirmed the frequent occurrence of multipolar spindles in two-cell stage *ssna-1(bs182)* embryos (6/14) (Pfister et al. 2024). Moreover, we generated *sas-1(t1476); ssna-1(bs182)* double-mutant embryos and found that, in contrast to either single mutant, sperm-contributed GFP::SAS-7 foci are not detected at pronuclear meeting (Fig. S5D), as is the case in the single *sas-1(is13)* null mutant (see Fig. 2H).

**Figure 5.**
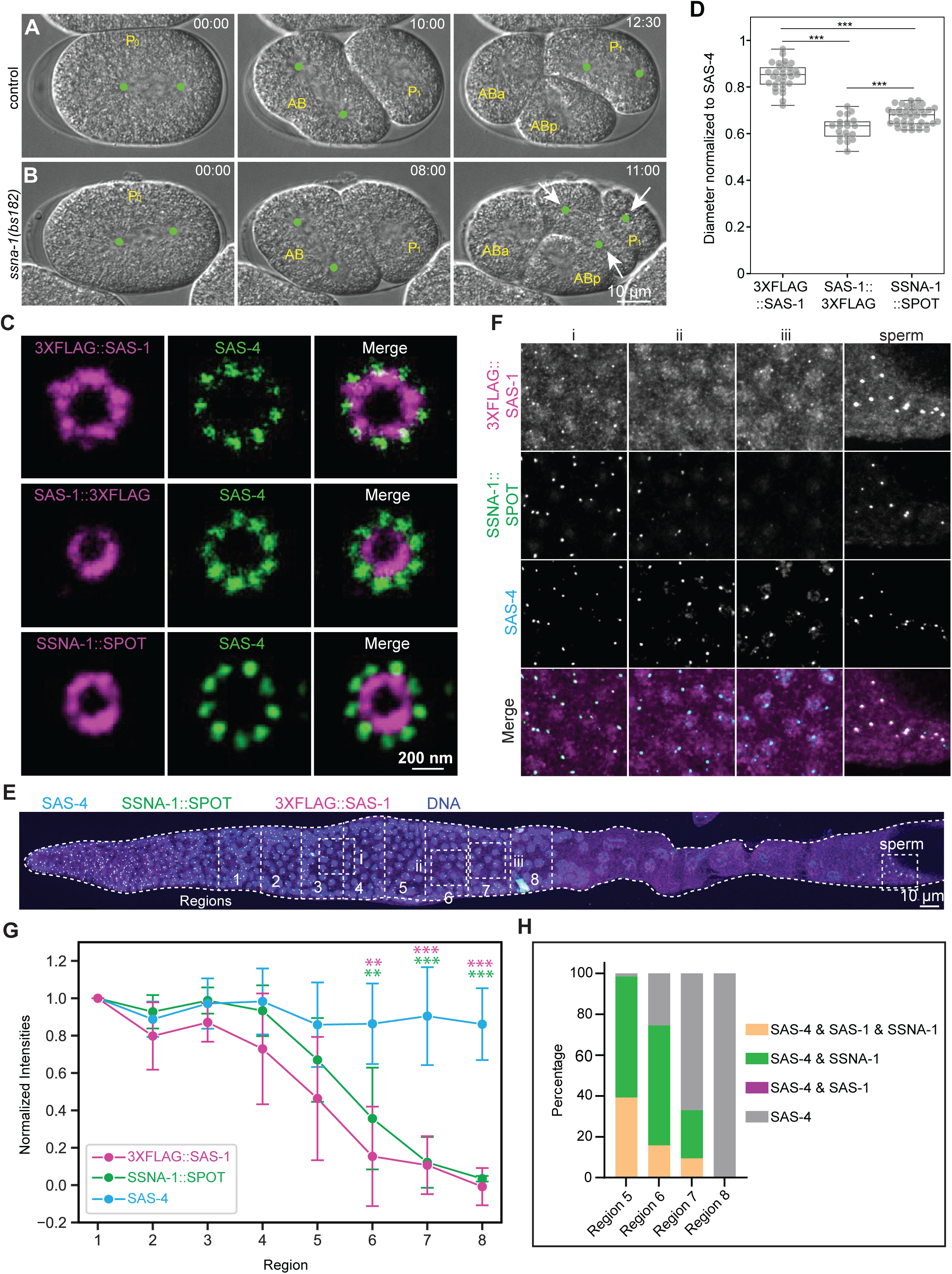
SSNA-1 is the part of central tube, together with the SAS-1 C-terminus. **A-B**. Snapshots from DIC live imaging of early embryos of control (A; N = 8) and *ssna-1(bs182)* embryo (B; N = 14). Time indicated in min:sec; images acquired every 30 sec. Cell designations are shown in yellow; green dots indicate spindle poles, arrows the poles of a tripolar spindle. **C.** Representative U-Ex-STED of meiotic prophase centrioles of animals expressing FLAG*-*tagged SAS-1 (on either N- or C-terminus, as indicated), and SSNA-1::SPOT, immunostained together with SAS-4. **D.** Quantification of diameter of 3XFLAG::SAS-1 (N = 31), SAS-1::3XFLAG (N = 20) and SSNA-1::SPOT (N = 40) distributions normalized with SAS-4 ring diameter. Student’s two-tailed t-tests, whereby *P* < 0.001 (***). **E.** Representative immunostaining of fixed gonads with antibodies against SAS-4 and FLAG, as well as SPOT. Different sections of gonad were stitched for visualization. The gonad is divided into eight regions, going backward from cellularization of meiotic nuclei (region 8). Numbered regions indicate where the quantification reported in G and H has been conducted. Dashed squares (i, ii, iii) indicate areas magnified in F. **F.** About three times magnified insets from (E) across meiotic prophase. **G.** Normalized signal intensities for SAS-4, 3XFLAG::SAS-1, and SSNA-1::SPOT across the eight regions indicated in (E). Background subtracted signal intensities for each region were averaged and then normalized with the mean intensity of region 1. Mean ± SD for each region (N= 6 gonads, with 15 – 25 foci per region for each gonad). Student’s two-tailed t-tests were performed for each region comparing signal intensities of SAS-4 with 3XFLAG::SAS-1 (magenta stars) and with SSNA-1::SPOT (green stars); *P* < 0.01 (**); *P* < 0.001 (***). **H.** Quantification of foci comprising SAS-1, SSNA-1 and SAS-4 in the indicated combination in the last four regions of (E) (N = 6 gonads; total number of foci analyzed: 197 for region 5, 133 for region 6, 106 for region 7, 74 for region 8).

SSNA-1 has been reported to localize inside the microtubule wall of centrioles (Pfister et al. 2024). We set out to compare the localization of SSNA-1 with that of SAS-1 using U-Ex-STED. The N-terminus of SAS-1 localizes next to centriolar microtubules and SAS-4 (Fig. 5C), compatible with this moiety of SAS-1 associating with microtubules, whereas the C-terminus maps to the more inward central tube (Fig. 5D) (Woglar et al. 2022). Importantly, using the diameter of the SAS-4 signal as a ruler in these experiments, we found SSNA-1::SPOT to co-localizes with the C-terminus of SAS-1 (Fig. 5D). Compatible with this view, AlphaFold2 multimer predictions suggest that SAS-1 interacts with SSNA-1 through its C-terminal alpha-helix (Fig. S5E, S5F), in line with structural evidence (Agostini et al. 2024). Overall, these findings suggest that SAS-1 and SSNA-1 somehow together contribute to centriole integrity.

### SAS-1 is required for SSNA-1 recruitment to centriole, cilia and microtubules

Given their close localization at centrioles and potential joint function, we addressed whether SSNA-1 departs from centrioles concomitantly with SAS-1 during oogenesis. As show in Figure 5E-H, we found that SSNA-1 leaves centrioles just after SAS-1, and before other centriolar components like SAS-4. Next, we investigated the epistatic relationship between SAS-1 and SSNA-1 during oogenesis. We found that TagRFP-T::SAS-1 departs precociously from centrioles in *ssna-1(bs182)* mutants (Fig. 6A-C). Strikingly, in addition, SSNA-1::SPOT is absent from centrioles in *sas-1(is13)* mutants (Fig. 6D). SSNA-1 also localizes at sensory cilia (Pfister et al. 2024), and we found a similar relationship between SAS-1 and SSNA-1 in that tissue, with SAS-1 levels being decreased at cilia in the absence of SSNA-1 (Fig. S6A, S6B) and SSNA-1 failing to localize at cilia upon neuron-specific SAS-1 depletion (Fig. 6E). Overall, we conclude that SAS-1 is essential for SSNA-1 localization at centrioles and at the ciliary base, whereas SSNA-1 reciprocally maintains SAS-1.

**Figure 6.**
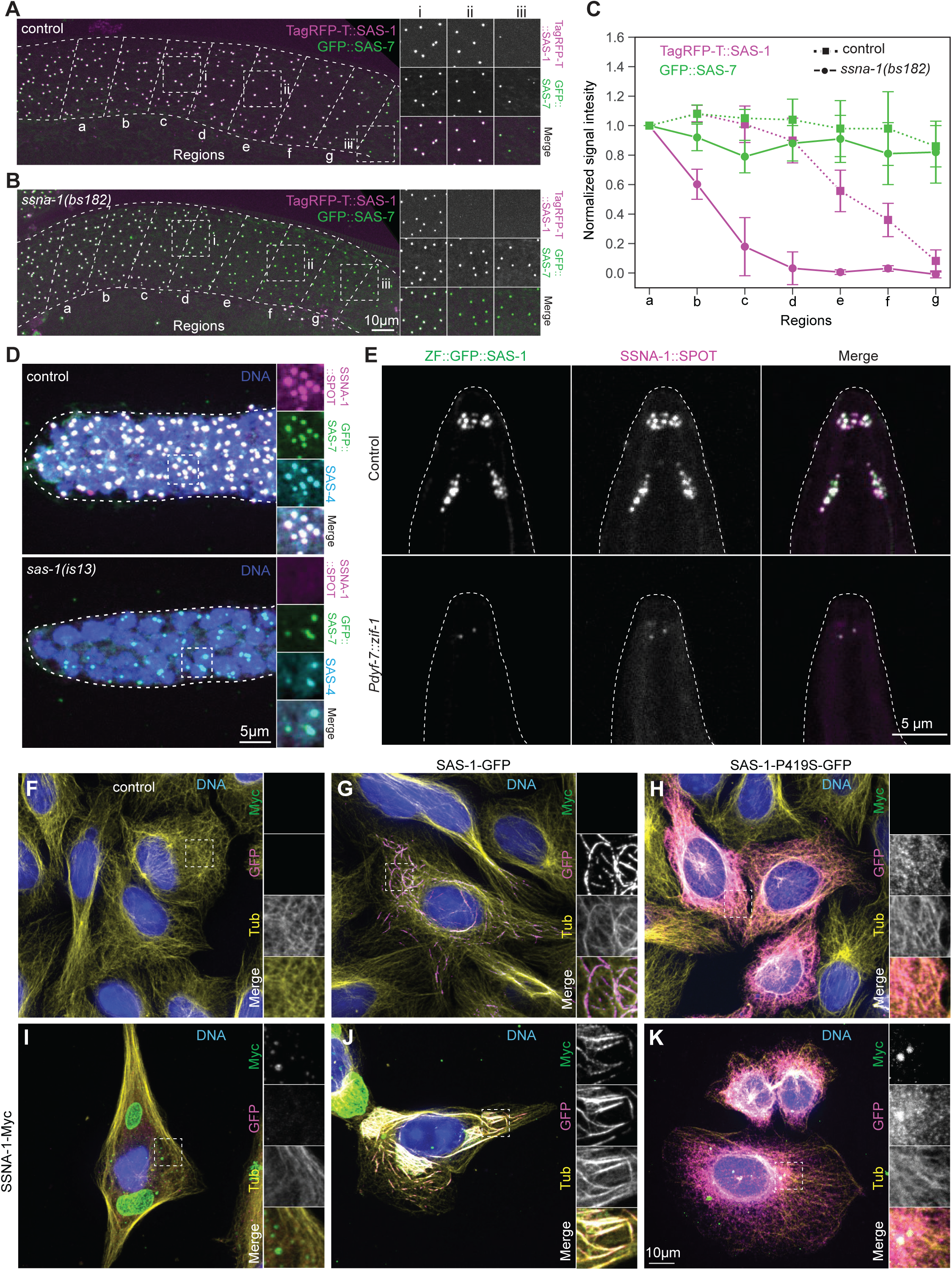
SAS-1 recruits SSNA-1 on centrioles and cilia. **A, B.** Live imaging of gonads in worms expressing GFP::SAS-7 and TagRFP-T::SAS-1 in control (A) and *ssna-1(bs182)* mutant (B). Dashed squares (i, ii, iii) indicate areas magnified approximately 1.5 times on the right. Gonads are divided into seven regions going backward from the loop region (g). **C.** Quantification of signal intensities for GFP::SAS-7 (green) and TagRFP-T:::SAS-1 (magenta) across regions indicated in (A) and (B). Background subtracted signal intensities for each region were averaged and then normalized with the mean intensity of region a. Mean ± SD for each region; N = 7 (control) and 9 (*ssna-1(bs182)*) gonads, with typically 5-10 foci quantified per region. **D.** Immunostaining of control (N = 6) and *sas-1(is13)* mutant (N = 8) gonads stained for SAS-4, GFP and SSNA-1::SPOT. Dashed squares indicate areas in the mitotic region of the gonad magnified approximately 1.5 times on the right. **E.** Representative immunofluorescence images of control L1 larvae (N = 10), as well as L1 larvae following ZIF-1-mediated SAS-1 depletion from ciliated neurons (N = 11), immunostained for SSNA-1::SPOT. **F-K.** Immunostaining of U2-OS cells expressing control (F, no transfection), SAS-1-GFP (G), SAS-1-P419S-GFP (H), SSNA-1-Myc (I), SAS-1-GFP and SSNA-1-Myc (J) or SAS-1-P419S-GFP and SSNA-1-Myc (K), immunostained for GFP (magenta), α-tubulin (yellow) and Myc (green). Boxes indicate location of insets, which are magnified approximately 2-fold.

To further elucidate the relationship between SAS-1 and SSNA-1, as well as to clarify their interaction with microtubules, we expressed the worm proteins in human U2-OS cells (Fig. 6F-K). Consistent with previous work (Von Tobel et al. 2014), we observed that wild-type SAS-1-GFP, but not a SAS-1-P419S-GFP protein corresponding to the *sas-1(t1476)* mutant, localizes with microtubules and induces their bundling (Fig. 6G, 6H). By contrast, SSNA-1-Myc forms large cytoplasmic aggregates that do not co-localize with microtubules (Fig. 6I). Importantly, in addition, we discovered that SAS-1-GFP co-expression recruits SSNA-1-Myc to microtubules (Fig. 6J). This colocalization is not observed when SSNA-1-Myc is co-expressed with SAS-1-P419S-GFP (Fig. 6K), together indicating that SSNA-1 localization on microtubules is contingent upon interaction with functional SAS-1.

## DISCUSSION

The mechanisms through which the centriole organelle maintains integrity once assembled are only beginning to be understood. Here, we analyzed how SAS-1 contributes to centriole integrity in *C. elegans*. Through the generation of a null allele and tissue-specific removal, we demonstrate that SAS-1 is dispensable for assembly of the centriole, but required for maintaining its integrity during oogenesis, spermatogenesis and embryogenesis. Moreover, we reveal that SAS-1 is present at the transition zone of sensory cilia and contributes to cilium function. Furthermore, our findings indicate that the ability of SAS-1 to bind microtubules enables recruitment of SSNA-1 to centrioles and cilia.

### SAS-1 is dispensable for centriole assembly but essential for centriole integrity

Centrioles are exceptionally stable organelles that are resistant to microtubule depolymerization induced by drugs or low temperature. Reflecting this unusual stability, several structural centriolar proteins exhibit at most marginal turnover once incorporated into the organelle. Thus, pioneering experiments in vertebrate cells established that centriolar α/ý-tubulin do not exhibit substantial exchange during an entire cell cycle (Kochanski & Borisy 1990). Likewise, in *C. elegans*, centriolar ý-tubulin, SAS-6 and SAS-4 incorporated in sperm centrioles undergo little to no exchange over many cell cycles in the ensuing embryo (Balestra et al. 2015). What then triggers programmed disassembly of such a stable organelle?

Previous analysis conducted with two reduction of function alleles, *sas-1*(*t1476*) and *sas-1*(*t1521*), indicated that SAS-1 is key for centriole integrity in *C. elegans* (Von Tobel et al. 2014; Pierron et al. 2023). However, because these are not null alleles, these early findings left open the possibility that SAS-1 is required for centriole assembly as well. To address this possibility, we generated the null allele *sas-1*(*is13*). Importantly, we found that centrioles are assembled during the proliferative cell cycles in the mitotic region of the gonad in *sas-1*(*is13*) mutants, as evidenced by the radial distribution of SAS-4 and microtubules, as well as by their capacity to seed the formation of procentrioles. However, these centrioles are slightly narrower and exhibit altered integrity, since SAS-4 and microtubules are compromised when analyzed by U-Ex-STED, likely reflecting their fragility.

In principle, the ability to undergo centriole assembly in the mitotic region of *sas-1*(*is13*) mutants could reflect unusually long perdurance of the maternal contribution. However, centrioles are also assembled, but not maintained, in *sas-1*(*is13*) mutant embryos, further attesting to a function strictly in the maintenance of centriole integrity. Therefore, the function of SAS-1 is distinct from that of the centriolar assembly factors SAS-7, SPD-2, ZYG-1, SAS-4, SAS-5 or SAS-6, without which centrioles do not assemble (reviewed in Ohta et al. 2017). Instead, we conclude that SAS-1 exerts a specific function in ensuring centriole integrity.

Components critical for centriole integrity have been identified in other systems, including human cells, where the inner scaffold that comprises notably POC1A/POC1B and WDR90 plays an important role (Steib et al. 2020; Sala et al. 2024). , 8- and χ-tubulin, as well as their interacting proteins TEDC1 and TEDC2, which are required for the assembly of microtubule doublets and triplets, are important for centriole integrity in human cells (Wang, Kong, et al. 2017; Breslow et al. 2018; Pudlowski et al. 2025). Upon their depletion, only microtubule singlets are present, and centrioles loose structural integrity during mitosis. The ciliopathy protein HYLS1 is also required for doublet and triplet microtubule formation, as well as for centriole stability (Takeda et al. 2024). Given that nematode centrioles harbor 9-fold radially symmetrical microtubule singlets, instead of the usual doublets and triplets, this explanation is unlikely to hold in *C. elegans*, although a B-like extension unveiled by cryo-electron tomography in centrioles of the embryo might contribute to stability (Tollervey et al. 2024). Regardless, SAS-1 plays a critical role for centriole integrity, and it will be interesting to test whether the SAS-1 homologue C2CD3 acts in a related manner.

### SAS-1 function at sensory cilia

We discovered a novel localization of SAS-1 at the transition zone of sensory cilia. Echoing the redundancy manifested by other components contributing to ciliary function in *C. elegans* (Williams et al. 2008; Schouteden et al. 2015), we found that SAS-1 contributes to ciliary function in a partially redundant manner with the transition zone protein NPHP-4. Given its localization within the confines of the axonemal microtubules, we speculate that SAS-1 may be part of the central cylinder, which also comprises CCEP-290 (Schouteden et al. 2015). By analogy with how SAS-1 operates in the context of centriole integrity, we further speculate that it may contribute to maintaining the ciliary axoneme in place during ciliogenesis. In human cells, C2CD3 is required for full-fledged centriole formation, as well as cilium generation, together with its upstream regulator Cep120 (Tsai et al. 2019). Therefore, in both worms and human beings, SAS-1/C2CD3 is critical for building a proper sensory cilium.

### Relationship of SAS-1 with SSNA-1

Although many proteins that reside at the centriole and the PCM have been identified in *C. elegans* through forward genetics and RNAi-based functional genomic approaches, together with others (Pfister et al. 2024), we report SSNA-1 as a novel centriolar component in the worm. In mammalian systems, SSNA1 has been described to localize to centrioles, primary cilia, mirroring the findings in worms, as well as axon branch sites (Goyal et al. 2014; Pfannenschmid et al. 2003; Pfister et al. 2024; Lawrence et al. 2021). *In vitro*, SSNA1 self-assembles into head-to-tail fibrils that have been reported to bind microtubules (Basnet et al. 2018). Likewise, *C. elegans* SSNA-1 can form such tetrameric polymer assemblies (Agostini et al. 2024). Furthermore, human SSNA1 has also been found to interact with the microtubule severing enzyme Spastin (Goyal et al. 2014).

Just like SAS-1, SSNA-1 contributes to centriole integrity in *C. elegans*, although to a lesser extent, judging from the milder phenotypes of *ssna-1*(*bs182*) null mutant animals (Pfister et al. 2024; this work). Accordingly, whereas SSNA-1 contributes to SAS-1 recruitment or maintenance at centrioles, we found that SAS-1 is essential for the presence of SSNA-1 at centrioles and cilia. Interestingly, recent work demonstrates that C2CD3 likewise recruits SSNA-1 to centrioles in human cells, mirroring the relationship between the related *C. elegans* proteins (Jen-Hsuan Wei, personal communication). This work also suggests that SSNA1 may not bind microtubules directly, as initially suggested (Basnet et al. 2018). In line with this suggestion, we found through AlphaFold2 modeling that the C-terminus of SAS-1 could interact with the SSNA-1 tetrameric polymer, whereas the N-terminus of SAS-1 is known to reside in the vicinity of the centriolar microtubules (Woglar et al. 2022), away from where SSNA-1 localizes. Furthermore, we found that *C. elegans* SSNA-1 does not associate with microtubules in a heterologous human cell assay, but is recruited to the polymer upon co-expression of functional SAS-1. Taken together, these findings indicate that SAS-1 is a microtubule-binding protein that, in conjunction with SSNA-1, forms the central tube of *C. elegans* centrioles. Furthermore, C2CD3 and SSNA1 appear to exhibit an analogous relationship in human cells (Jen-Hsuan Wei, personal communication). Given that SSNA1 interacts with the microtubule severing enzyme Spastin in human cells (Goyal et al. 2014), perhaps C2CD3/SAS-1 together with SSNA1/SSNA-1 shield centriolar microtubules from being severed precociously.

## Conclusion

In conclusion, our analysis with novel deletion and tissue-specific depletion alleles of SAS-1 sheds new light on the mechanisms governing integrity of the centriole organelle.

## ACKNOWLEDGMENTS

We thank Kevin O’Connell and Jason Pfister for sharing *ssna-1*(*bs182*) *and ssna-1::spot* worm strains prior to publication, as well as for fruitful discussions, Jen-Hsuan Wei for communicating results prior to publication, Cédric Pourroy for help with the SPY650-tubulin ciliary marking experiment, Alana Dastous for help with AlphaFold multimer, Rémi Dornier (BioImaging and Optics Platform of the School of Life Sciences, EPFL) for a macro to quantify GFP::SAS-1 foci. We are grateful to Friso Douma, Gabriela Garcia-Rodriguez and Marie Pierron for constructive comments on the manuscript. This work was supported by the Swiss National Science Foundation (grant # 310030_197749).

## MATERIALS AND METHODS

### C. elegans strains

Worms were grown using standard protocols on nematode growth medium (NGM) plates seeded with *Escherichia coli* OP50 as food source (Brenner 1974). Strains used in this study are listed in Table S1. Animals were typically raised at 20-22°C except for temperature-sensitive strains which were grown at 16°C until the L4 stage and then shifted to 24°C before imaging in adulthood. Synchronized worm populations were obtained either by bleaching gravid worms and letting embryos hatch in M9 minimal medium, or by allowing 20-30 gravid adult worms to lay embryos on NGM plates for 1-2 hours and then removing the adults.

### Cell culture

U2-OS cells were cultured at 37°C with 5% CO2 in DMEM supplemented with GlutaMAX (ThermoFisher Scientific) and 10% fetal bovine serum. For the U2-OS-SAS-1-GFP and U2-OS-SAS-1(P419S)-GFP stable cell lines (Von Tobel et al., 2014), the medium was supplemented with 1 µg/mL puromycin. For transient expression experiments, full-length *ssna-1* cDNA fused to a Myc tag was cloned into a pDONR221 vector using BP Clonase reaction (ThermoFisher Scientific). Next, *ssna-1* was transferred to a pEBTet destination vector using LR Clonase reaction, where *ssna-1* is under the control of a doxycycline-inducible CMV promoter. Cells were seeded on coverslips in a 6-well plate and allowed to grow to ∼60-70% confluency before transfection, which was performed using Lipofectamine 3000 (ThermoFisher Scientific) following the manufacturer’s protocol using 2 µg of the cloned *ssna-1*-Myc plasmid. Post-transfection, cells were incubated for 4-6 hours, after which the medium was replaced with fresh medium containing 1 µg/ml doxycycline (Merck). Cells were then allowed to express the transfected construct for 48 hours, before fixation with prechilled methanol at -20°C for 7 min, followed by three washes with PBS and immunostaining.

### Gonad and sperm spreading

Gonad spreading was performed similarly to Woglar et al. 2022. In brief, gonads from 1000-2000 young adult hermaphrodites were dissected in 30 μL of PBS (20% in H2O) with Levamisole (1 mg/mL). 10 μL of this solution was put on a clean 22 x 40 mm coverslip and 50 μL spreading buffer [32 μL of fixation solution (4% w/v Paraformaldehyde and 3.2% w/v Sucrose in H2O), 6 μL of Lipsol solution [1% v/v Lipsol in H2O] and 2 μL of Sarcosyl solution (1% w/v of Sarcosyl in H2O)] was added. The coverslips were dried for 1 hour at room temperature and a further 3 hours at 37°C. Thereafter, coverslips were either stored at -80°C or processed further for immunostaining or expansion microscopy. For sperm spreading, ∼1000 hermaphrodites or males were washed and resuspended in 100 µL 1x nuclear purification buffer (10 mM HEPES, 40 mM NaCl, 90 mM KCl, 2 mM EDTA, 0.5 mM EGTA, 0.2 mM DTT, 0.5 mM Spermidine, 0.1% Triton-X). The worms were then homogenized by twenty strokes with a tight pestle with a 90° turn at each stroke, before spinning down 30 μL of this suspension onto 8 mm round coverslips (10,000g for 5 min). The coverslips were removed and the specimen fixed with prechilled methanol at -20°C for 5 min, washed further with PBST and processed further for expansion microscopy.

### Gonad and sperm expansion for U-Ex-STED

Ultrastructure expansion microscopy coupled to STED was performed based on Woglar et al. 2022. Briefly, dried coverslips containing spread gonads were further fixed with prechilled methanol at -20°C for 20 min, followed by three 10 min washes with PBS-T (this step is skipped for spread sperm nuclei). The coverslips were then incubated overnight at room temperature in Acrylamide/Formaldehyde solution (1%/0.7% in PBS). On the following day, the coverslips underwent three 5 min washes with PBS. For the gelation step, 22 x 40 mm coverslip were incubated with ∼250 μL (20 μL for 8 mm coverslips) of monomer solution (19% (wt/wt) Sodium Acrylate, 10% (wt/wt) Acrylamide, 0.05% (wt/wt) BIS in PBS), supplemented with 0.5% Tetramethylethylenediamine (TEMED) and 0.5% Ammonium Persulfate (APS) for 1 hour in a humid chamber protected from light. The resulting gels were then incubated with denaturation buffer (200 mM SDS, 200 mM NaCl, and 50 mM Tris (pH 9)) for 1 hour at 70°C. Post-denaturation, gels were washed five times with distilled water for 20 min each, followed by an overnight incubation in distilled water. The expansion factor was calculated by measuring the dimensions of the expanded gels with a ruler and comparing them with those of the coverslips. For immunostaining, the gels were first incubated in blocking buffer (10 mM HEPES (pH 7.4), 3% BSA, 0.1% Tween 20, 0.05% sodium azide) for 1 hour at room temperature. They were then incubated overnight with primary antibodies diluted in blocking buffer at room temperature. The next day, after three 10 min washes with blocking buffer, gels were incubated with secondary antibodies and 0.7 μg/L Hoechst 33258 for 3 hours at 37°C. Post-incubation, gels were re-expanded through three 10 min incubations in distilled water. Finally, gels were mounted between two 22 x 60 mm coverslips coated with poly-D-lysine (2 mg/ml in water). The mounted gels were sealed using VaLaP, a 1:1:1 mixture of petroleum jelly, lanolin, and paraffin wax.

### Immunostainings

Embryo immunostaining was performed based on (Gönczy et al. 1999). Embryos were dissected from gravid hermaphrodites in 6 µL of water on Poly-D-Lysine (2 mg/ml in water) coated frosted slides (Marienfeld, 1000200). An 18 x 18 mm coverslip was placed gently on the embryos and excess water removed with a blotting paper, leading to slight flattening of the embryos. The slide was quickly placed on a prechilled metal block kept on dry ice for a minimum of 10 min. Thereafter, the coverslip was flicked using a scalpel or razor blade and the slide dipped in prechilled methanol at -20°C for 2 min for fixation. Slides were then washed 3 times in PBST to remove traces of methanol, followed by blocking for 30 min in 2% BSA in PBST. Primary antibodies in blocking buffer were incubated either for 1-2 hours at room temperature or overnight at 4°C. Slides were then washed three times in PBST for 10 min each. Secondary antibodies in PBST were incubated at room temperature for 45 min, followed by washing and 0.7 μg/L Hoechst 33258 staining in PBST. The slides were washed twice before mounting in 4% n-Propyl-Gallate, 90% Glycerol, 1x PBS, and coverslips sealed with colorless nail polish.

Immunostaining on whole dissected gonads was performed based on Pierron et al, 2023. Gonads from young adult hermaphrodites were dissected in sperm buffer (50 mM HEPES [pH 7.0], 50 mM NaCl, 25 mM KCl, 5 mM CaCl2, 1 mM MgSO4, 50 mM Glucose, 1 mg/mL BSA) with Levamisole (1mg/mL), and gently placed on frosted glass slides coated with Poly-D-Lysine (2 mg/mL in PBS), followed by lowering of the coverslips (22 x 40 mm) and freezing on dry ice. The coverslip was flicked as above and the slide dipped in prechilled methanol at -20°C for 5 min. The remaining steps, including washing, blocking and antibody incubation, were similar to those mentioned for embryo immunostaining.

For immunostaining of U2-0S cells, fixed cells were blocked for 30 min using 3% BSA in with 0.05%(v/v) Tween20 (PBST), followed by primary antibody incubation in the blocking buffer for 1 hour at room temperature. This was followed by three washes with PBST, and secondary antibody incubation with 0.7 μg/L Hoechst 33258 for 1 hour at room temperature. Cells were washed again, stained with 0.7 µg/L Hoechst 33258 for 5 minutes, and mounted on slides using Fluoromount G (Invitrogen).

Primary antibodies used in this study: rabbit anti-SAS-4: 1:5000 (immunostaining or U-Ex-STED)(Leidel & Gönczy 2003); rabbit anti-SAS-6: 1:800 (immunostaining) or 1:500 (U-Ex-STED) (Leidel et al., 2005); rabbit anti-GFP: 1:1000 (immunostaining) (gift from Viesturs Simanis); rabbit anti-α-tubulin 1:5000 (Western blotting) (Abcam | ab52866); mouse anti-FLAG: 1:500 (U-Ex-STED) (Thermo | MA1-91878); mouse anti-FLAG: 1:200 (immunostaining), 1:3000 (Western blotting) (Merck | F1804); SPOT: 1:500 (U-Ex-STED) (Chromotek | 28a5-20); mouse anti-α/β-tubulin: 1:500 (immunostaining and U-Ex-STED) (Lima & Cosson 2019); mouse anti-Myc 1:500 (immunostaining) (CST | Ab#2276S); SPOT nanobody Alexa Fluor 568: 1:1000 (immunostaining) (Chromotek | ebAF568); SPOT nanobody Alexa Fluor 488: 1:1000 (immunostaining) (Chromotek | ebAF488).

The following secondary antibodies were used at 1:1000 (immunostaining) or 1:500 (U-Ex-STED): goat anti-mouse Alexa Fluor 488; goat anti-rabbit Alexa Fluor 568; donkey anti-rabbit Alexa Fluor 594; goat anti-mouse Alexa Fluor 647.

### Microscopy

Three confocal microscopy systems were utilized. First, an upright Leica SP8 equipped with two hybrid photon counting detectors (HyD) and a transmission photomultiplier tube (PMT) for brightfield imaging. This system employs a 63x HC Plan-Apochromat objective (NA 1.4) and 405, 488, and 552 nm solid-state laser lines for excitation, paired with a DFC 7000 GT (B/W) camera. Second, an inverted Olympus IX83 motorized microscope equipped with Yokogawa spinning disk CSU-W1 head, a 60x (NA 1.42 U PLAN S APO) objective, as well as ImagEMX2 EMCCD and Orca Flash 4.0 sCMOS cameras, with image acquisition controlled by VisiView software. Furthermore, 2D-STED (Stimulated Emission Depletion) images were captured on a Leica TCS SP8 STED 3X microscope, using a 100x 1.4 NA oil-immersion objective. The system employs 488 and 589 nm excitation lasers, along with 592 and 775 nm pulsed lasers for depletion.

Live imaging of late-stage *C. elegans* embryos (Fig. 3A) was performed using a Zeiss Lattice Lightsheet 7 microscope equipped with a 44.83×/1.0 NA water-immersion objective. Embryos were maintained at 22–24 °C during imaging. Volumetric image stacks were acquired every 10 minutes using 488 and 561 nm solid-state lasers for excitation; images were captured on a PCO Edge 4.2 sCMOS grayscale camera.

### Image processing and analysis

Visualization and quantification of microscopic images were performed using Fiji (ImageJ) (Schindelin et al. 2012). For quantification of nuclear area in Fig. 1E, the Z-plane containing the largest cross-sectional area of each nucleus was identified. An ellipse was visually fitted to this plane, its area measured and used as a proxy for nuclear size.

For quantification of 2D-STED images (Fig. 1G-I and Fig. 2B-E), individual channels were manually aligned based on centriole ring structures to correct for chromatic shifts. A 1-pixel Gaussian blur was applied to all images for both analysis and display. For display purposes only, to account for variable zoom factors across image sets, regions of interest (ROIs) of identical pixel dimensions were drawn on each image and then uniformly scaled to match the smallest field of view. Brightness and contrast were individually adjusted for each image. To measure centriole diameter, a line was drawn across each centriole, and a super-Gaussian curve fitted to the intensity profile. The full width at half maximum (FWHM) of the fit was taken as the centriole diameter.

For Fig. 3B, GFP::SAS-1 foci were quantified from Lattice lightsheet microscopy datasets using a custom Fiji macro. A user-defined ROI was drawn to restrict analysis to the embryo region, and a Laplacian of Gaussian (LoG) detector was applied frame-by-frame to identify potential centrioles. Spot detection was performed on a per-channel and per-timepoint basis, using subpixel localization and intensity thresholds optimized for each channel. Only foci within the user-specified ROI were retained, and their positions and intensities were saved for further analysis.

For measurements of cilia and dendrite length (Fig. 4L-Q and Fig. S4C), segmented lines were drawn on maximum-projected images. Dendrite length was measured from the neuron cell body to the base of the cilium, cilium length from the ciliary base to the tip of the axoneme.

For signal intensity quantification in Fig. 5G and Fig. 6C, sum-projected images of gonads were divided into eight (Fig. 5G) or seven (Fig. 6C) regions defined by ∼3× the diameter of diplotene nuclei - measured at the onset of cellularization (Fig. 5G) or at the gonad loop (Fig. 6C). ROIs of 1.8 μm² were placed on each focus and an adjacent background area. Background-corrected intensities were calculated per channel and normalized to the most distal region. For Fig. S2D-I, 15 × 15 pixel ROIs were centered on each focus and a nearby background region at the z-slice of maximal intensity. A 6.3 µm sum projection centered on this slice was generated, and background-subtracted signal was measured. For Fig. S6A, ROIs of ∼12 µm² were drawn around amphid cilia and adjacent background regions on a ∼7 µm sum projection. Background-subtracted intensities were reported.

### Ciliary assays

For cilia staining, L1-L2 worms were washed in M9 buffer and incubated with 100 nM SPY650-tubulin (Spirochrome) for 3 hours. After incubation, worms were washed again in M9 buffer and immediately imaged.

The chemotaxis assay was conducted based on Bargmann et al. 1993. Approximately 50–100 well-fed adult worms were collected from NGM plates and placed in an Eppendorf tube with M9 buffer, and washed twice with M9 to remove any residual bacteria. A 10 cm NGM assay plate was prepared by marking two points in a straight line along its diameter and located near the periphery of the plate. At one of these points, 1 µL of 100% isoamyl alcohol (attractant) mixed with 1 µL 10% sodium azide was applied. At the opposite point, 1 µL of 100% ethanol (repellent) mixed with 1 µL 10% sodium azide was added. The sodium azide served to paralyze the worms, facilitating accurate counting. The washed worms were then transferred to the center of the plate, approximately 5 cm from both attractant and repellent point sources. After 1 hour, the distribution of worms was analyzed. Worms that had migrated toward either end - i.e., within approximately a 2.5 cm radius of the attractant or repellent point were counted and analyzed.

The chemotaxis index was calculated using the following formula:

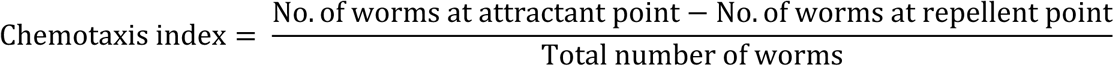

The chemotaxis index ranged from -1 (perfect repellent) to 1 (perfect attractant).

---

**Table S1:** list of strains used in this study.

| Lab designation | Genotype | References |
| --- | --- | --- |
| GZ1385 | *sas-7(or1940[gfp::sas-7])III; glo-1(zu931)X* | (Sugioka et al. 2017) |
| GZ2134 | *sas-7(or1940[gfp::sas-7])III; sas-6(ok2554)IV/nT1(qls51)IV; glo-1(zu931)X* | (Sugioka et al. 2017; Kitagawa et al. 2011); this study |
| GZ2122 | *sas-7(or1940[gfp::sas-7])III sas-1(is13)III/hT2; ssna-1(bs206 [ssna-1::spot]) IV* | (Sugioka et al. 2017; Pfister et al. 2024); this study |
| GZ1816 | *sas-4(bs195[sas-4::gfp] III* | (Kalbfuss & Gönczy 2023a) |
| GZ2140 | *sas-4(bs195[sas-4::gfp] III sas-1(is13)III/hT2; ssna-1(bs206 [ssna-1::spot]) IV* | (Kalbfuss & Gönczy 2023a; Pfister et al. 2024); this study |
| GZ2123 | *zif-1(gk117) III sas-7(is1[tagRFP::sas-7+loxP])III sas-1(is12[zf::gfp:sas-1]III); ssna-1(bs206 [ssna-1::spot]) IV* | (Klinkert et al. 2019; Pfister et al. 2024); this study |
| GZ1754 | *stIs10544 [hlh-16::H1-wCherry::let-858 3' UTR], sas-7(or1940[gfp::sas-7])III, glo-1(zu931)X* | (Kalbfuss & Gönczy 2023a) |
| GZ2135 | *stIs10544 [hlh-16::H1-wCherry::let-858 3' UTR], zif-1(gk117) III sas-7(is1[tagRFP::sas-7+loxP])III sas-1(is12[zf::gfp:sas-1]III)* | (Kalbfuss & Gönczy 2023a); this study |
| GZ2050 | *sas-7(or1940[gfp::sas-7])III sas-1(is10[TagRFP-T::sas-1])III; glo-1(zu931)X* | (Sugioka et al. 2017); this study |
| GZ2071 | *spd-5(vie26[gfp::spd-5 +loxP])I; sas-1(is10[TagRFP-T::sas-1])III* | (Pierron et al. 2023); this study |
| GZ2118 | *vieSi22[pAD401; Pmksr-2::gfp::mksr-2; cb unc-119(+)]I; sas-1(is10[TagRFP-T::sas-1])III; glo-1(zu931)X* | (Schouteden et al. 2015); this study |
| GZ2115 | *zif-1(gk117)III sas-7(is1[tagRFP::sas-7+loxP])III sas-1(is12[zf::gfp:sas-1]III); vieSi70[pAD675;pche-11::che-11::mKate2] IV* | (Klinkert et al. 2019; Garbrecht et al. 2021); this study |
| GZ2106 | *ItSi1017[pDC335; Pdyf-7::vhhgfp4::ZIF-1;cb-unc-119(+)] II; unc-119(ed3) III? zif-1(gk117)III sas-7(is1[tagRFP::sas-7+loxP])III sas-1(is12[zf::gfp:sas-1]III); vieSi70[pAD675;pche-11::che-11::mKate2] IV* | (Klinkert et al. 2019; Garbrecht et al. 2021); this study |
| GZ2119 | *zif-1(gk117)III sas-7(is1[tagRFP::sas-7+loxP])III sas-1(is12[zf::gfp:sas-1]III); vieSi70[pAD675;pche-11::che-11::mKate2] IV; nphp-4(tm925)V* | (Schouteden et al. 2015; Klinkert et al. 2019; Garbrecht et al. 2021); this study |
| GZ2120 | *ItSi1017[pDC335; Pdyf-7::vhhgfp4::ZIF-1;cb-unc-119(+)] II; unc-119(ed3) III? zif-1(gk117)III sas-7(is1[tagRFP::sas-7+loxP])III sas-1(is12[zf::gfp:sas-1]III); vieSi70[pAD675;pche-11::che-11::mKate2] IV; nphp-4(tm925)V* | (Klinkert et al. 2019; Garbrecht et al. 2021); this study |
| GZ2116 | *sas-7(or1940[gfp::sas-7])III sas-1(is10[TagRFP-T::sas-1])III; ssna-1(bs182)/ears-2(ve631[LoxP + myo-2p::GFP::unc-54 3' UTR + rps-27p::neoR::unc-54 3' UTR + LoxP]) IV* | (Sugioka et al. 2017; Pfister et al. 2024); this study |
| GZ2080 | *sas-1(is7[3xflag::sas-1])III; ssna-1(bs206 [ssna-1::spot]) IV* | (Woglar et al. 2022; Pfister et al. 2024) |
| GZ1914 | *sas-7(or1940[gfp::sas-7])III, sas-1(t1476)/hT2; glo-1(zu931)X* | (Von Tobel et al. 2014; Sugioka et al. 2017); this study |
| GZ2141 | *sas-7(or1940[gfp::sas-7])III sas-1(t1476)/hT2 III; ssna-1(bs182)/ears-2(ve631[LoxP + myo-2p::GFP::unc-54 3' UTR + rps-27p::neoR::unc-54 3' UTR + LoxP]) IV* | (Von Tobel et al. 2014; Sugioka et al. 2017; Pfister et al. 2024); this study |
| GZ2076 | *sas-7(or1940[gfp::sas-7])III sas-1(is10[TagRFP-T::sas-1])III; ssna-1(bs182)/ears-2(ve631[LoxP + myo-2p::GFP::unc-54 3' UTR + rps-27p::neoR::unc-54 3' UTR + LoxP]) IV* | (Von Tobel et al. 2014; Sugioka et al. 2017; Pfister et al. 2024); this study |
| GZ2136 | *ItSi1017[pDC335; Pdyf-7::vhhgfp4::ZIF-1;cb-unc-119(+)] II; unc-119(ed3) III? zif-1(gk117)III sas-7(is1[tagRFP::sas-7+loxP])III sas-1(is12[zf::gfp:sas-1])III); ssna-1(bs206 [ssna-1::spot]) IV* | (Klinkert et al. 2019; Garbrecht et al. 2021; Pfister et al. 2024); this study |

**Figure S1:**
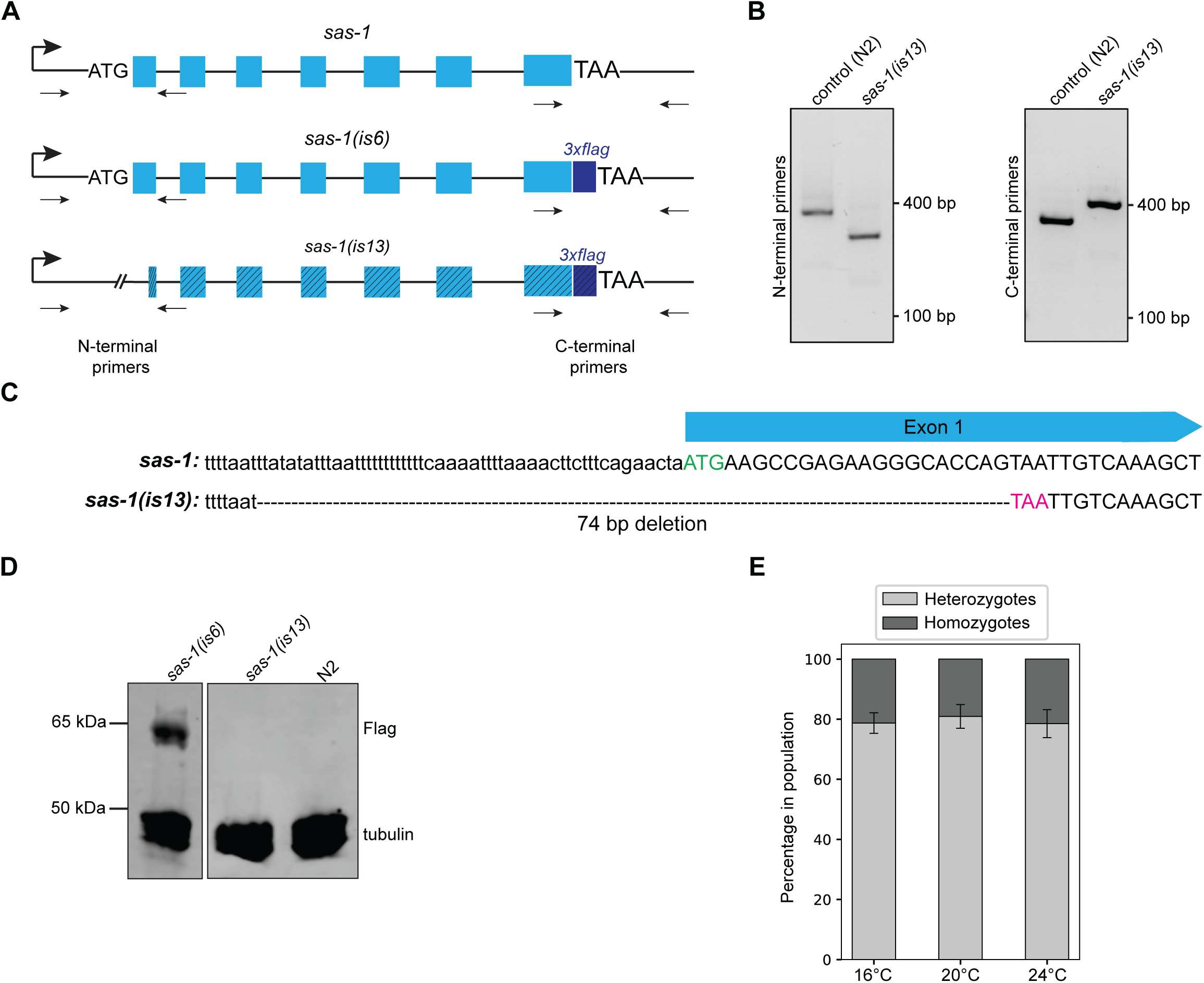
Generation of *sas-1* null allele. **A.** Schematic of wild-type *sas-1*, as well as *sas-1(is6)* (Woglar et al. 2022), and *sas-1(is13)* mutant alleles. Primers at the N- and C-terminus used for PCR-based genotyping are indicated. **B.** Agarose gel image of PCR products from genomic DNA of control and *sas-1(is13)* mutant worms using primers mentioned in (A). **C.** Sanger sequencing revealed a 74 bp deletion in *sas-1(is13)*, stretching from the 5’-UTR to Exon 1, leading to loss of the ATG start codon. **D.** Western blotting of worm lysates. Note that the ∼65 kDa band of *sas-1::3xflag* is lost in *sas-1(is13)*. **E.** Progeny test of *sas-1(is13)/hT2(gfp)* worms; N = 8 technical repeats.

**Figure S2:**
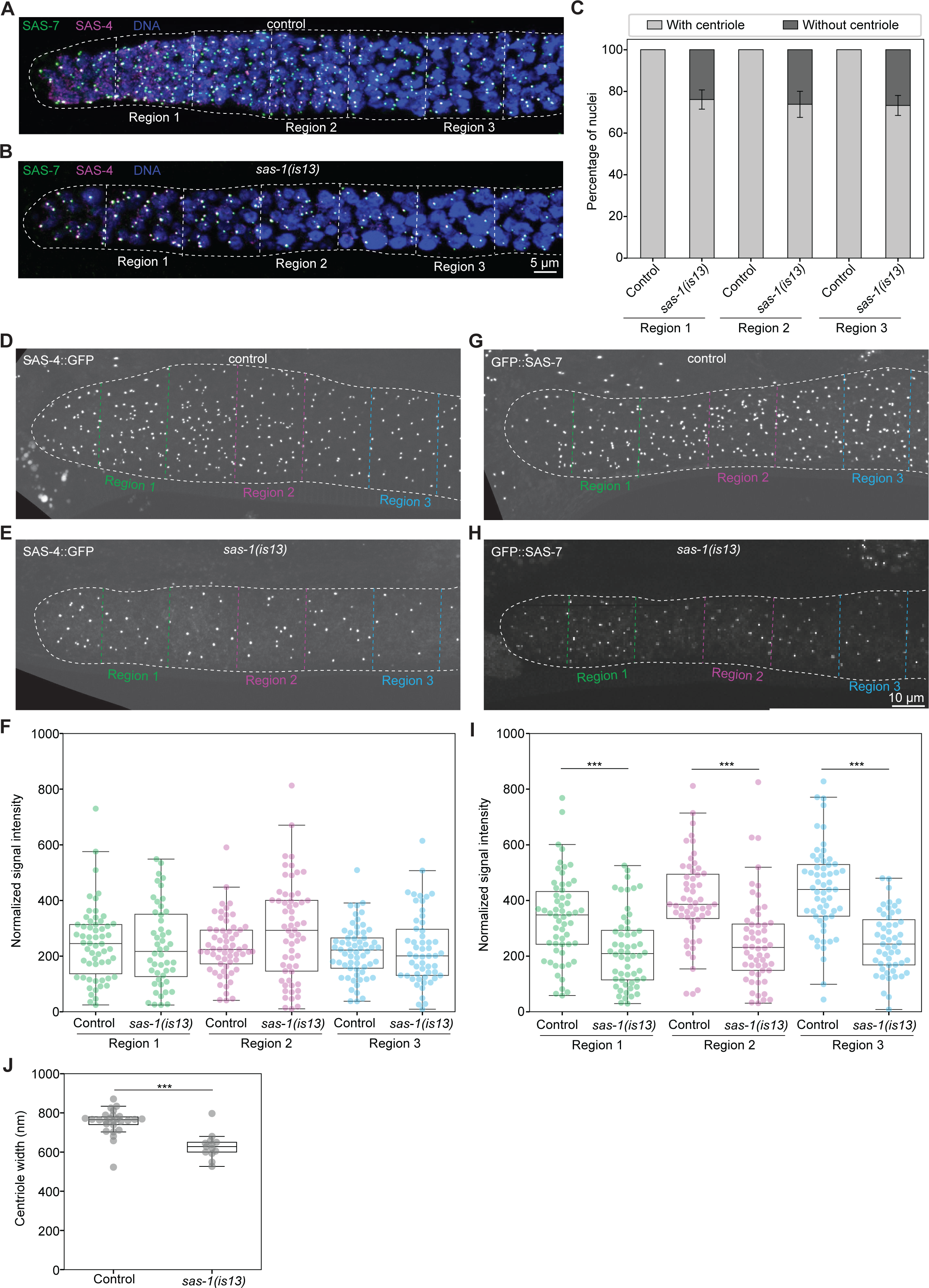
Level of SAS-7 and SAS-4 in *sas-1(is13)* **A, B.** Representative immunofluorescence images of gonads of control (A) and *sas-1(is13)* mutant (B), both expressing GFP::SAS-7 and immunostained for GFP and SAS-4. **C.** Quantification of percentage of nuclei with or without centrioles in different regions of distal gonad of control (N *=* 6 gonads) and *sas-1(is13)* mutants (5 gonads; N = 230 for region 1; N = 236 for region 2; N = 208 for region 3). **D, E, G, H.** Live imaging of mitotic region of the gonad in control (D, G) or *sas-1(is13)* mutant (E, H) worms expressing SAS-4::GFP (D, E) or GFP::SAS-7 (G, H). Three regions of about 18 µm in width and 18 µm apart were defined as indicated. Signal intensity of SAS-4::GFP and GFP::SAS-7 was then quantified in these regions. **F, I.** Quantification of signal intensity of SAS-4::GFP (F) and GFP::SAS-7 (I) in the three regions described in (D, E, G, H). Foci number for F: control (5 gonads; N *=* 58 for region 1, 61 for region 2, 61 for region 3), *sas-1(is13)* (5 gonads; N *=* 48 for region 1, 60 for region 2, 54 for region 3). Foci number for I: control (5 gonads; N *=* 58 for region 1, 55 for region 2, 59 for region 3), *sas-1(is13)* (5 gonads, N *=* 56 for region 1, 56 for region 2, 51 for region 3). Student’s two-tailed t-tests, whereby *P* < 0.001 (***). The differences between control and *sas-1*(*is13*) in F are not significant (Region 1: *P* = 0.77, Region 2: *P* = 0.055, Region 3: *P* = 0.69). **J.** Quantification of centriole width in the mitotic region of control (N = 26) and *sas-1(is13)* mutants (N = 13), excluding centriole fragments. Student’s two-tailed t-tests, whereby *P* < 0.001 (***).

**Figure S3:**
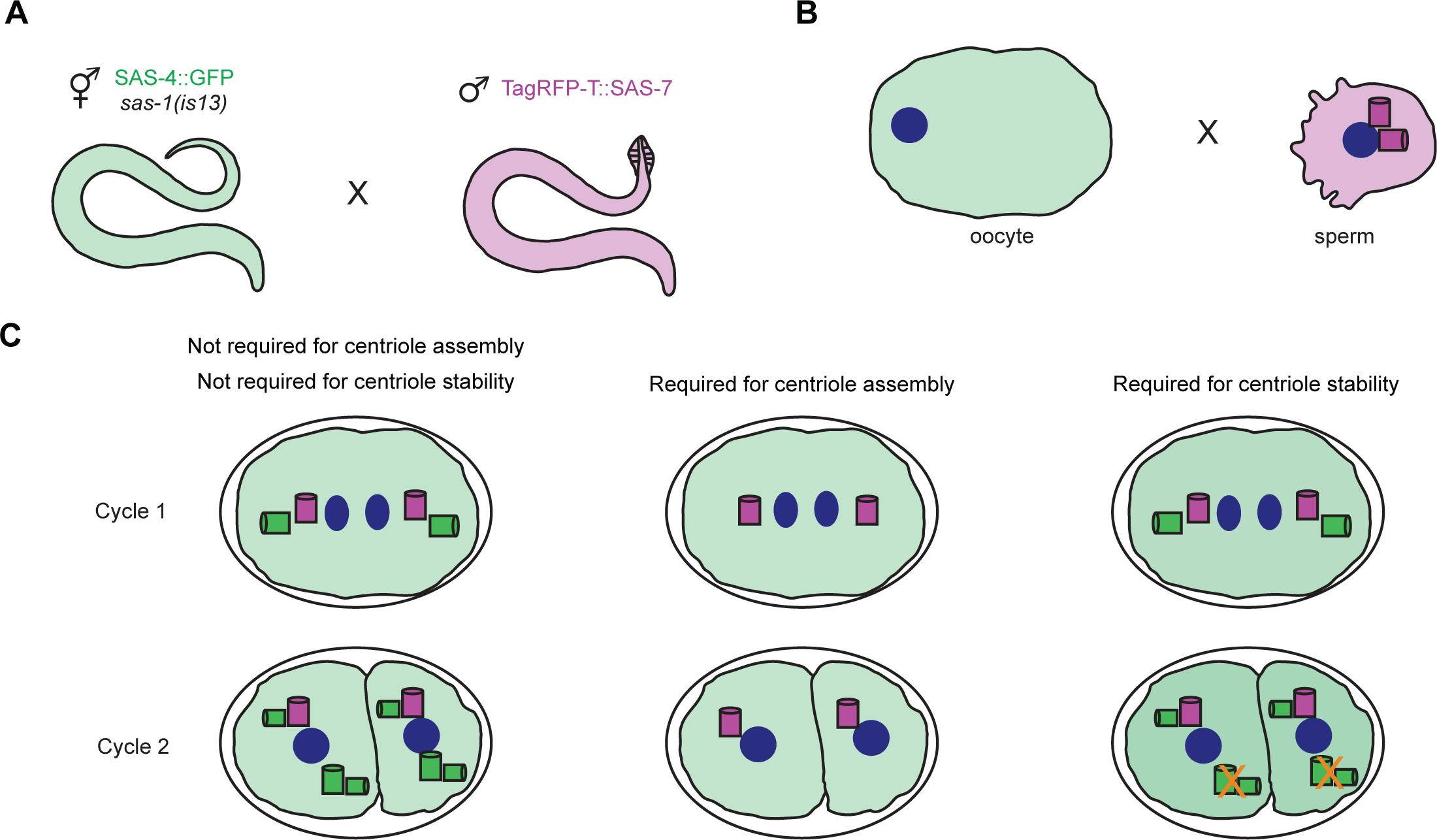
Schematic of marked mating experiment. **A, B.** *sas-1(is13)* mutant hermaphrodites expressing SAS-4::GFP (green) mated with control males expressing TagRFP-T::SAS-7 (magenta). **C.** Possible outcomes of marked mating experiment: (left) SAS-1 is not essential for centriole assembly or stability; in this case, in cycle 2, two SAS-4::GFP foci are present in both blastomeres, which undergo bipolar spindle assembly. (middle) SAS-1 is essential for centriole assembly; in this case, no SAS-4::GFP foci is present in the embryo, and monopolar spindle assembly occurs in both blastomeres at the two-cell stage. (right) SAS-1 is essential for centriole stability; in this case, two SAS-4::GFP foci are present in each blastomere initially, but are eliminated thereafter (indicated by orange cross).

**Figure S4:**
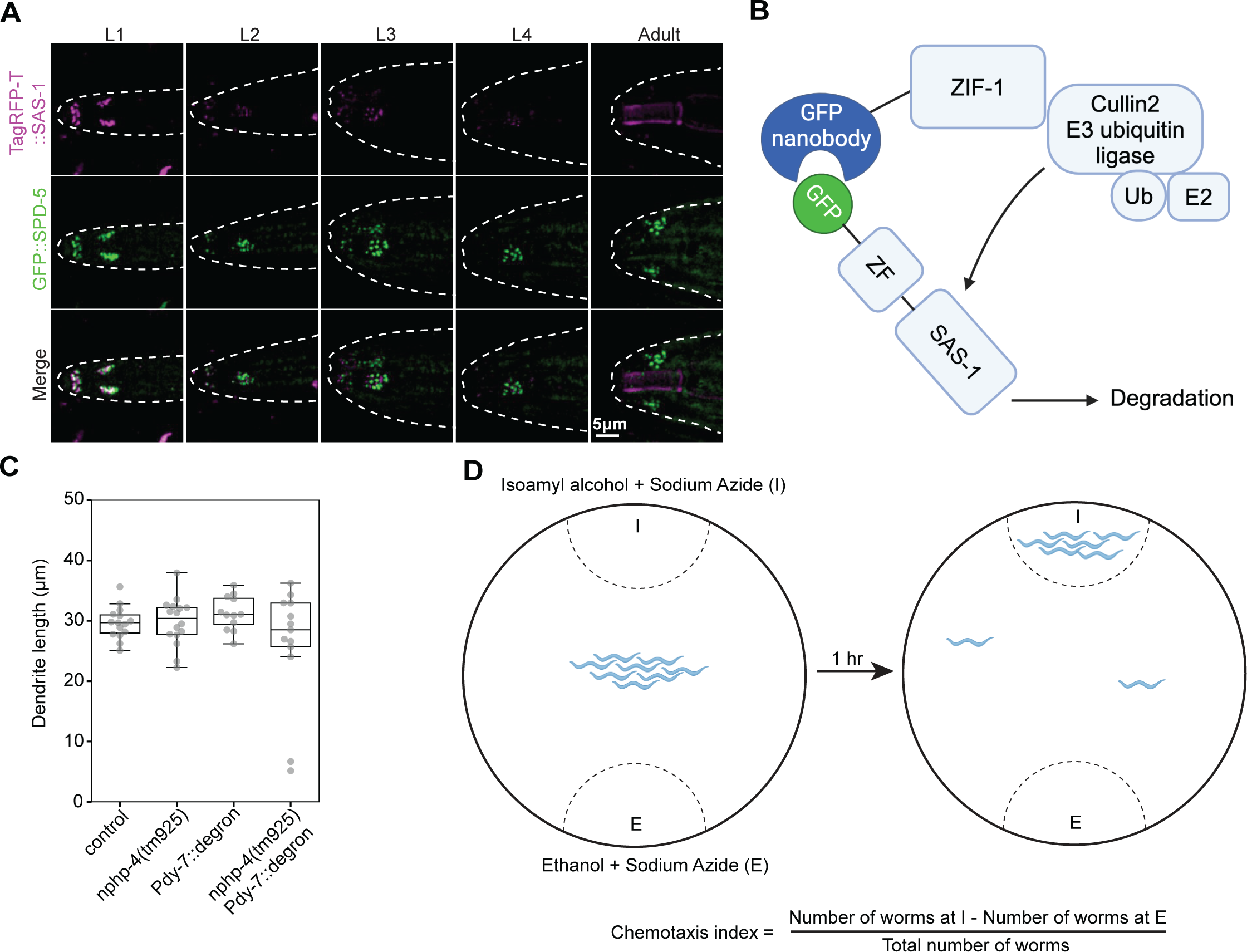
SAS-1 plays a role in cilia function. **A.** Live imaging of sensory cilia at anterior of animals expressing TagRFP-T::SAS-1 and GFP::SPD-5 during larval stages and adulthood, as indicated. N *=* 10 for each stage. The bright cylindrical signal observed in the TagRFP-T channel in the adult stage is likely autofluorescence from the mouth orifice. **B.** Schematic of ZIF-1 mediated degron system, with ZIF-1 expression under the control of the *dyf-7* promoter in this case. **C.** Quantification of dendrite length in Fig. 4L-P. Animals quantified: N *=* 15 (control); 16 (*nphp-4(tm925)*, 12 (*Pdyf-7::degron*), 13 (*Pdyf-7::degron; nphp-4(tm925))*. Student’s two-tailed t-tests, which were all not significant compared to control (*nphp-4(tm925)*: *P* = 0.88, *Pdyf-7::degron: P* = 0.142, *Pdyf-7::degron; nphp-4(tm925), P* = 0.19). **D.** Schematic of chemosensation assay (not to scale) (drawn with Biorender). See text for further details.

**Figure S5:**
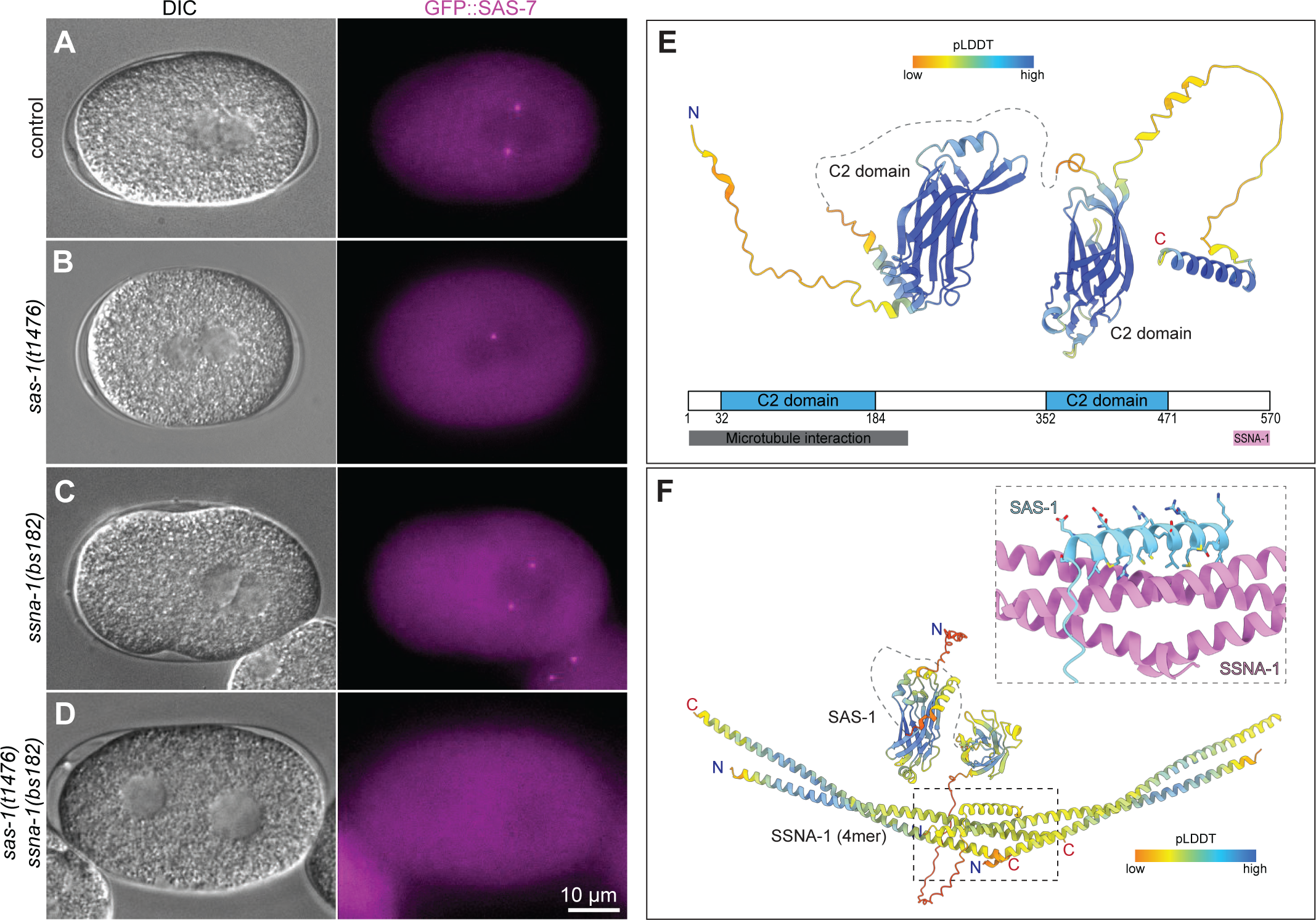
Interaction between SAS-1 and SSNA-1. **A-D.** Snapshots from time-lapse imaging showing DIC and GFP::SAS-7 distribution in embryos at pronuclear meeting in control (A; N = 10), *sas-1(t1476)* (B; N = 10), *ssna-1(bs182)* (C, N = 10), and *sas-1(t1476); ssna-1(bs182)* (D; N = 7) embryos. DIC channel is a single plane whereas the GFP::SAS-7 channel is a maximum intensity projection of selected Z-planes. **E.** AlphaFold2 prediction of SAS-1 with two C2-domains and a C-terminal alpha helix. The flexible loop connecting the two C2 domains is indicated with dotted line. **F.** AlphaFold2 prediction of SAS-1 binding to a SSNA-1 tetramer. SSNA-1 is forming a head-to-tail fibril with the C-terminal helix of SAS-1 binding to the interface of two helixes. Top-right: magnified representation of the interface marked with dotted line; SAS-1 in cyan, SSNA-1 in pink.

**Figure S6:**
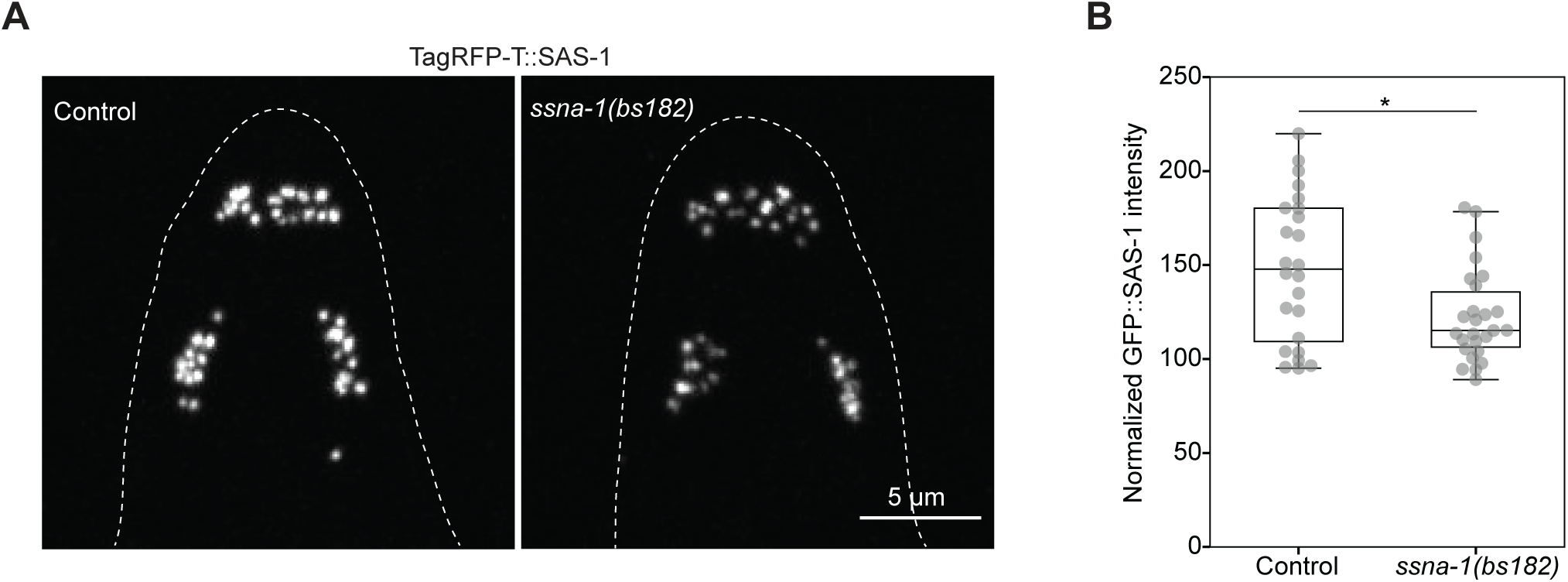
SAS-1 contributes to SSNA-1 distribution at sensory cilia. **A.** Live imaging of anterior cilia of L1 larvae expressing TagRFP-T::SAS-1 in control (N *=* 12) and *ssna-1(bs182)* mutant (N *=* 13). **B.** Quantification of TagRFP-T::SAS-1 signal intensity of anterior sensory cilia in control (N =24) and *ssna-1(bs182)* mutant (N = 26) L1 larvae. Student’s two-tailed t-tests, whereby *P* < 0.05 (*).

